# Golden section hypothesis of macroevolution: unification of metabolic scaling and *Fibonacci* sequence

**DOI:** 10.1101/2022.09.19.508476

**Authors:** Xin G. Yang, Lei Wang

**Author notes:** Author for correspondence: Xin G. Yang.

## Abstract

Golden section is a subtle technology from nature to split space, which is both extensive and mysterious. In recent years, some studies1-4 have begun to focus on metabolic scaling (*B*∝*M^b^*) at the macroevolutionary scale, and some important trends have been revealed. To further answer the question of "where does b come from and where does it go in evolution", a golden section model of macroevolution was constructed by integrating metabolic scaling and Fibonacci sequence. The results showed that, (1) macroevolution at the boundary level was a highly ordered process from one-dimensional (prokaryotes) to five-dimensional evolution (fungi). Four-dimensional life^5^ was only the choice of animals. (2) b just was the syndrome of dimension application and metabolism realization of life following Fibonacci sequence; however, it indicated major evolution events in the macroevolution and the directions in secondary macroevolution. The logic and panorama of macroevolution therefore were re-outlined based on the idea of dimensional evolution and metabolic evolution. It was argued that the golden section model of macroevolution established a full-new logic system of dimensional and metabolic evolution, and provided a possible path for the unification of macroevolution and microevolution.

---

Scaling relations offer powerful insights into the fundamental processes that constrain and regulate biological structure and function^2, 6^. Metabolic scaling was a relationship function between individual metabolic rate (*B*) and individual mass (*M*), which was generally expressed as *B*∝*M^b^*. Kleiber’s Law^7^, *B*∝*M*^3/4^ was the most famous one. For a long time, scientists have tried to unify the metabolic scaling theory based on Kleiber’s law. The early work of metabolic scaling focused on the unified model of *b* across species^8–13^.

In recent years, when there has been increasing evidence supporting the nonuniqueness of *b* cross species, research has begun to focus on the integration of macroevolution and metabolic scaling^1–4^, especially with the gradual improvement of the large database of cross-species metabolic scaling. Relevant studies have found that there were indeed some regular changes in metabolic scaling in macroevolution, such as the allometric scaling convergence hypothesis^2^ and metabolic homeostasis hypothesis^1^. Article^4^ published in *PNAS* in 2019 fully refreshed researchers’ understanding of the pattern of metabolic scaling in macroevolution, based on one of the world’s largest integrated databases of metabolic scaling. However, these studies based on statistical laws still cannot answer a fundamental question: where does *b* come from, and where does it go in evolution?

This paper put forward a macroevolutionary assumption combined with the latest progress of metabolic scaling studies and taking the results of several major databases of metabolic scaling in macroevolution as the test^1–4^: *Macroevolution was the evolutionary arrangement of life forms corresponding to a series of optimized choices between growth and metabolism, along the realization track of maximizing metabolic efficiency (i.e. the values of b).* The following questions were: (1) was there a necessary connection between the occurring sequence of life forms and the size distribution of the *b* value (i.e. where does *b* go) in macroevolution? (2)Was there a definite metabolic scaling theory or other theoretical basis for this connection (i.e. where does *b* come from)? It seemed that a rule needed to be found that was superior to the current statistical law of metabolic scaling according to the systematics principle, not just the experience summary and theoretical induction based on the statistical law, to answer the two questions.

The golden section was a universal law in nature, that spanned biology and physics, and there was a definite mathematical function, namely, the Fibonacci sequence^14^. Figuratively speaking, golden section was a subtle space segmentation (*f*(*n*)) with time as scissors (*n*), and a natural dichotomy for the optimal use of space. One famous case of a golden section applied in biology was Ludwig’s law. In the existing metabolic scaling theory, the WBE model^8^ and Yoda self-thinning rule^15^ were typical representatives of the optimal utilization of biological space, which revealed the biological space growth mode of animal individuals and plant populations for optimization of metabolic efficiency respectively. Therefore, the golden section was essentially unified with the WBE model and Yoda self-thinning rule.

Based on the above analysis, this paper proposed and verified a golden section model of macroevolution integrating metabolic scaling and Fibonacci sequence, by a technical route of priori setting - logical analysis - geometric derivation - comparative verification. There were three grounds for doing so. First, cell growth, as the underlying mechanism of metabolic scaling, and Fibonacci series were both exponential functions based on the natural logarithm *e*, which were unified in mathematical form. Second, both the evolution trajectory of life and the Fibonacci function were so called inherited expansions based on some dimensions, which were unified in function properties. Third, both metabolic scaling (such as the WBE model and Yoda model) and the golden section were related to the biophysical law of optimal space utilization, which was unified in the model mechanism.

## Hypothesis

### Priori model

A priori model was set such that there was the following relationship in macroevolution,

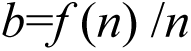

*b*, Characteristic value of the metabolic scaling function (*B*∝*M^b^*). *f*(*n*) and *n* were from *Fibonacci* functions, which satisfy the following functional relationship,

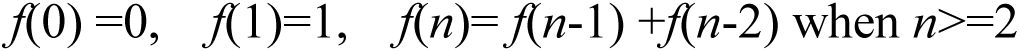

The value range of *n* was 1, 2, 3, 4, 5, which successively corresponded to prokaryotes, protists, plants, animals and fungi. The meanings of *n* and *f*(*n*) in macroevolution are also limited here. First, *M^f^*^(*n*)^ ∝*B^n^* can be derived based on the priori model. Therefore, *n* in macroevolution reflected an eigenvalue of growth efficiency based on metabolism (*B*). Similarly, *f*(*n*) was a metabolic eigenvalue based on growth (*M*).

As shown in table 1, there seemed to be a metabolic evolutionary cycle from prokaryotes (*b*=1) to fungi (*b*=1) in macroevolution. 3/4 of animals and 2/3 of plants were basically in line with the existing knowledge, and 1(5/5) of fungi were also in line with general experience, despite the lack of scaling research in fungi. However, the results of prokaryotes and protists were not completely consistent with the existing metabolic scaling database^1, 2, 4, 16^.

**Table 1.** *b* values derived by Fibonacci sequence in macroevolution.

| Basic variables | life form at the boundary level |  |  |  |  |  |
| --- | --- | --- | --- | --- | --- | --- |
|  | - | Prokaryotes | Protists | Plants | Animals | Fungi |
| $n$ | 0 | 1 | 2 | 3 | 4 | 5 |
| $f(n)$ | 0 | 1 | 1 | 2 | 3 | 5 |
| $b$ ($=f(n)/n$) | - | 1 | 1/2 | 2/3 | 3/4 | 1 |

### Premise setting in the derivation of priori model

First, more derivation was paid to the generality of *b* in macroevolution at the boundary level, rather than the diversity of *b* in secondary macroevolution (such as phylum, or class). However, based on the principle of systematic evolution, higher levels of macroevolution had general binding effect on the lower.

Second, a unified paradigm of “Optimized growth mode in biological space” was constructed through decomposition before integration of two spatial growth modes, i.e. *D*_B_ (metabolic dimension, spatial geometric growth of "functional components" that determine individual metabolic efficiency) and *D*_M_ (growth dimension, spatial geometric growth of individual itself). And *b*=*D_B_*/*D_M_* was set according to the biological dimension idea of metabolic scaling^5^. As shown in table 2, there might be five space growth modes defined by *D_M_* from prokaryotes to fungi in turn, according to the priori model.

**Table 2.** Optimized growth mode in biological space in macroevolution.

| Life forms | Variables |  |  |  |
| --- | --- | --- | --- | --- |
| | $b$ | $D_B$ | $D_M$ | space growth modes |
| **Prokaryotes** | 1 | 1 | 1 | One-dimensional |
| **Protists** | 1/2 | 1 | 2 | Two-dimensional |
| **Plants** | 2/3 | 2 | 3 | Three-dimensional |
| **Animals** | 3/4 | 3 | 4 | Four-dimensional |
| **Fungi** | 1 | 5 | 5 | Five-dimensional |

Third, complete metabolic paths generally include absorption, transportation, transformation and so on. There were one or more key metabolic links to determine the optimization of metabolic efficiency^17^, and became the basis of “Optimized growth mode in biological space”. Similarly, the resource supply may also become the basis^17^.

## Derivation and verification

### Prokaryotes b=f (1) /1=1/1=1, D_B_=1, D_M_=1

#### Origin of one-dimensional growth of prokaryotes in macroevolution

From the evolutionary perspective of spatial growth mode, prokaryotes (such as bacteria), as unicellular organisms, mostly existed in groups. However, it was only a cell cluster even if they gathered together, which was undoubtedly the most unfavorable for the spatial utilization of each organism. For example, Staphylococcus had a population growth pattern that reduced the cost of aggregation, through the axis-direction elongation of the cluster. Therefore, one-dimensional elongation for the cells was undoubtedly a reasonable choice of spatial growth mode in evolution to expand the scope of spatial search. This was called one-dimensional growth. From sphaerita, and bacilli to spirochetes (including actinomycetes), this obvious evolutionary trend within prokaryotes still can be seen. At the same time, this one-dimensional growth was also conducive to maximizing the relative surface area of the cell (i.e., the ratio of surface area to volume), to obtain greater metabolic efficiency. For the prokaryotes with one-dimensional growth (geometric derivation can be found in methods), there was *A∝L∝L^D_B_^*, *V∝L∝L^D_m_^*, therefore, *D*_B_=1, *D*_M_=1, and *b*=1/1=1. *L,* biological length, *A,* biological area, *V,* biological volume.

In addition, prokaryotes have innate conditions and accompanying evolutionary skills to adapt to one-dimensional growth. First, the requirement of metabolic activities on the spatial coordination of multiple metabolic sites in cells was lower, since for their imperfect cell function and structure (Theoretically, spherical cells made it easier to achieve spatial coordination of inner functions). Second, it was the evolution of multinucleated cells, which further overcame the influence on metabolism from the increase of internal distance in cells. At the same time, the increases in genetic material in multinucleated cells also led to significant increases in metabolic efficiency^2^.

One-dimensional growth was a typical strategy of dimension reduction in evolution, compared with normal cells. Prokaryotes can achieve higher metabolic efficiency through one-dimensional growth, but that would also bring some problems to their overall development. In fact, the main body of prokaryotes remains is composed of bacilli, rather than spirochetes. It was argued that was a selective balance between dimension application and metabolism achievement in prokaryotes.

#### Comparison of b between empirical values and model derived value

It was easy to infer that *b*=2/3 for sphaerita, *b*=1 for spirochetes, and bacilli in the middle or close to *b*=1, based on similar analysis methods. The range of *b* values of prokaryotes should be 2/3∼1. Of course, the upper limit of the *b* value would be significantly greater than 1 if the potential impact of genetic material was considered^2^.

Prokaryotes were the group with the largest change in *b* values, and with the worst convergence of data distribution among all groups, based on results from a few major databases^1, 4, 16^. The measured values of *b* tended to exceed 1, which was called supermetabolic scaling^2^. Obviously, this was related to the diversity of dimension application at the taxonomic level and multinucleate cell evolution. Another reason might be that the metabolic homeostasis mechanism^1^ has not been established.

However, the *b* value for prokaryotes derived from the priori model was not the maximum value pursued by metabolic evolution, but the metabolic characteristic value indicating the one-dimensional spatial growth mode. This was a problem that needed to be distinguished in discussing the maximization of metabolic efficiency at different evolutionary scales.

#### Evolutionary significance of one-dimensional growth

However, the evolutionary logic of the one-dimensional growth of prokaryotes in macroevolution still existed. In contrast, it was the one-dimensional growth that just laid the structural foundation of the supermetabolic scaling^2^. Since it would be easier to realize the fusion of genetic material through the winding contact of linear cells, and then increase cell genome sizes^18^. At the same time, it was speculated that the one-dimensional growth certainly played key roles in the subsequent evolution, especially for the mycelial route closely related to the evolution of fungi.

### Protists b=f (2)/2=1/2, D_B_=1, D_M_=2

#### Origin of the two-dimensional growth of protists in macroevolution

Studies showed that the geometric shapes of individuals (such as plates, spheres and cylinders) can significantly affect the metabolic scaling^19^. The protist continued the metabolic pursuit by maximizing the relative surface area of the cell. There were two ways to achieve this, one was to reduce the cell volume, and the other was to increase the cell surface area. Obviously, early protists chose the latter^20^. The result of morphological evolution brought by this growth mode was the overall flattening and irregularity of cells, such as diatoms, amoebas, and myxomycetes. Geometrically, it led to the individual growth dimension *D*_M_ approaching two, namely, two-dimensional growth. At the same time, this two-dimensional growth also promoted the transfer of the key link limiting metabolic efficiency from absorption (determined by relative surface area of cell) to others since the area had been increased enough. Particularly, the spatial coordination of multiple metabolic sites within the cell would be more critical with the increasing of cell sizes^2^, relatively which would never arise in prokaryotes. Thus, in a flattened growing cell, the distance along its axis ultimately became the primary limiting parameter of metabolic efficiency, and the geometric attribute of this distance determined the metabolic dimension *D_B_*=1. Therefore (geometric derivation can be found in methods), *b*= *D_B_*/ *D_M_* =1/2.

#### Comparison of b between empirical values and model derived value

The statistical convergence of the *b* value for protists was greatly improved compared with prokaryotes, which was highly consistent with a trajectory line of *b*=1, similar to that of all species (from bacteria to animals), based on the results of a few major databases^1, 4^. An early study^16^ also showed that the empirical range of *b* values of protists was 2/3∼1. Therefore, the actual *b* value of protists in the current taxonomic system must be significantly higher than 1/2, and might show the statistics law of isometric metabolism scaling^2^.

This paper also made a geometric derivation of the *b* value of protists in current taxonomic system. The result was *b*=3/3=1 for Chlorella and *b*=4/4=1 for paramecium (Appendix 2). These results were highly consistent with the above empirical results^1, 2, 4^ and those of other species in protest^21, 22^, even similar to very small animals and plants^23, 24^. Therefore, the *b* value should be 1/2∼1, if early protists (such as diatoms, amoebas, slime bacteria, etc.) and the latter were combined. However, *b*=1/2 only appeared or applied to early protists. Similar to prokaryotes, the *b* value derived from the priori model was not the maximum value of metabolic evolutionary selection, but the metabolic optimization eigenvalue that indicated the two-dimensional growth.

#### Evolutionary significance of two-dimensional growth

Formally, the contradiction between dimension application and metabolism achievement in protists was the most prominent in macroevolution. However, while the metabolic efficiency decreased for the protists with two-dimensional growth, it also represented a significant increase in the growth efficiency (*M*∝*B*^2^). Evidence was obtained from the work of Payne^20^. Fossil evidence showed the first sudden increase in individual size occurring in the protist stage in the over 3.5 billion years of evolution. This also revealed another potential huge evolutionary advantage brought by the two-dimensional growth: the significant increase in cell size created more space for the further evolution of cell function.

Thus, it can be inferred that in the evolution stage of unicellular life, the differentiating selection between growth (by two-dimensional growth) and metabolism (by one-dimensional growth) had begun, and the significant fluctuation of metabolic efficiency^1, 2, 4^ in the stage therefore was not unrelated to this. The solution of contradiction appeared after the emergence of multicellular eukaryotes, and the individual size entered the second significant increasing stage^20^.

Protists, mainly including algae, protozoa, and protomycetes, are generally classified as the most primitive and simplest kind of unicellular eukaryote and a kind of life form with the largest variation in morphology, anatomy, ecology and life history. Just as reflected by the current taxonomic system of protists, the two-dimensional growth in evolution should result in a high degree of diversification of cell morphology and function, and even hasten the evolution of eukaryotic cells. For example, the folding and internalization of cell membranes was more likely to occur in two-dimensional cells, rather than spheres. Even, it was speculated that early prokaryotes (or archaea) had both one-dimensional and two-dimensional branches in evolution, and the two-dimensional branches gave birth to early protists due to the acceleration of eukaryotic evolution (Fig.4). The direct evidence came from the fact that organisms with two-dimensional forms only did not exist among the remains of prokaryotes. Another piece of evidence^25^ came from molecular evolution which revealed the possible relationship between early protest and archaea, based on their common cell membrane remodelling abilities.

In any case, both the one-dimensional growth of prokaryotes and the two-dimensional growth of protest indicated some important evolutionary events in the early macroevolution stage, not just the sizes of *b*.

### Plants b=f (3)/3=2/3, D_B_=2, D_M_=3

#### Origin of three-dimensional growth in macroevolution

*b*=2/3 should be universal as long as the spatial growth mode strictly follows the Euclidean geometric principle^5, 13, 15, 26^. It was speculated that there was one evolutionary route maintaining normal cell morphology in the early stage (such as from Sphaerita to Chlorella) until plants. Molecular evidence^27^ also showed that the genetic distance between plants and animals (or fungi) was longer than that between fungi and animals, indicating that there may be an independent evolutionary route for plants. Compared with the one-dimensional and two-dimensional routes, it was called the three-dimensional conservative evolutionary route.

However, for early aquatic photosynthetic bacteria and unicellular photosynthetic algae, their photosynthetic metabolic organs were mainly chlorophyll macromolecules filled in the globular cells, rather than in the cell surface, to obtain higher metabolic rates (*b*=*D_B_/D_M_*=3/3=1 can be deduced when metabolic rate was limited by cell volume^13^, similar to *Saccha romyces cerevisiae*^21^ and *Polytoma papillatum*^22^), which was also an adaptation in evolution for those unicellular living in a water environment, when nutrients and the cell surface had not become the first limiting factors^2^.

Only after the leaves and vascular bundles were evolved and then the geometric spatial behavior of growth and metabolism was separated, higher terrestrial plants finally evolved to the 2/3 rule as the metabolic rate isometric to surface area^13^.

#### Dimensional evolution logic of three-dimensional growth

Dimensional evolution in macroevolution got back on track from the beginning of the plant. Multicellular structure endow plants with the rapid development of space utilization ability, especially with the emergence of leaves and vascular bundles. The differentiation of multicellular structure supported plants approaching the two-dimensional resource spaces (such as lights limited by its incidence mode) by the three-dimensional individual growth, a coevolution process of the growth dimension and metabolic dimension. Tall and mature trees with "bundle vascular" could be regarded as the top of space utilization ability following Euclidean space rules in evolution, and finally showed compliance with mechanical laws^28, 29^, *H*∝*D*^2/3^, *H*, tree height, *D*, trunk diameter. From the perspective of light utilization, plant evolution could be viewed as a wrestling and rising process between mechanical limiting and height competition. The relation between the mechanical law and metabolic scaling for tall and mature trees was also discussed in the paper, more content can be found in the methods and Appendix 3.

There have been many debates about whether the 3/4 rule is suitable for plants. In fact, even if the key metabolic links of higher vascular plants were transformed into material transport within the vascularture, which was still different from the general logic of metabolic dimension generation of animals^8, 30^. One case of 3/4 in plants is further discussed in Appendix 3. It was argued that higher vascular plants conformed to the traditional Euclidean geometric principle, that is, *A∝L*^2^*∝L^D_B_^*, *V∝L*^3^*∝L^D_M_^*, therefore, *D*_B_=2, *D*_M_=3, and *b*=2/3, whether in individual or group living states following energy equivalence^31^.

#### Comparison of b between empirical values and model derived value

It was speculated that the *b* value of plants varied from 2/3∼1. In the empirical results^1, 4, 32^, *b* of individual trees tended to be close to 2/3, or tended to the mechanical laws, *H*∝ *D*^2/3^. 3/4 may also appear in empirical research^9, 24^, and in fact, it can be used as the optimum empirical value in forest management, as the best fitting mean value of 2/3∼1. However, this did not mean that plants chose 3/4 as the general model of dimensional evolution in macroevolution. Similarly, the *b* value derived from the priori model was not the maximum value of metabolism for plants, but the metabolic optimization eigenvalue that indicated the three-dimensional growth.

### Animals b = f (4) /4 = 3/4, D_B_=3, D_M_=4

#### Comparing among existing metabolic scaling theory

The theoretical research on the 3/4 metabolic scaling was most extensive, and the theoretical prediction was also consistent with empirical results, especially for higher animals^8, 10, 11, 30^.

In those studies, the WBE model^8^ and its proposed idea of biological dimension evolution^5^ provided an excellent demonstration for us to understand life evolution from the perspective of biological spatial growth. However, it was argued that the least carrier network theory^30^ was more general for animals, which revealed the generality of spatial geometry in the generation of metabolic dimensions for animals. The geometric rule therefore changed^8, 30^ as *V∝L*^4^*∝M, A∝L*^3^*∝B*. Therefore, *b*=3/4.

More debate about the 3/4 rule focused on its uniqueness across species. However, increasing evidence^1–4^ has showed that 3/4 might only be a dimension application evolved by higher thermostatic animals, and even small animals and ectotherms did not meet. This problem can be easily found by comparing the empirical results obtained from databases of different sizes and different grouping criteria^2, 4^.

#### Origin of four-dimensional growth in macroevolution

A very important basis of dimensional evolution was the adaptation of life to the suppling pattern of resources^2, 13, 17, 33^. From prokaryotes to plants (including fungi), the optimization of the space growth mode just were based on the physical spatial characteristics of the resource supply. However, animals were the first to break through the limitation after the protozoon changed their feeding mode and evolved an internal circulation system. The space growth mode of animal metabolism could be vividly called "spatial internalization", meaning that three space dimensions directly were given to *D_B_*, which was completely interlinked with^30^ *B*∝*L*^D^, *D* (=3) was the physical space dimensions occupied by animal bodies. In other words, the evolution direction had been doomed just from the moment that paramecium opened its mouth.

From this point of view, "spatial internalization" was also interlinked with the resource limitation mechanism in the quantum metabolic model^34^: the metabolic scaling followed the 3/4 rule for organisms subject to ecological constraints defined by *scarce but reliable resources* (for example, animals); the 2/3 rule was followed when ecological constraints were defined by *sufficient but only temporarily available resources* (for example, plants).

Simultaneously, animals must have an additional growth dimension to carry this internalized three-dimensional metabolic space. According to the metabolic optimization principle and the law of mass conservation^30^, the additional dimension could not be greater than 1 in evolution; otherwise, the metabolic efficiency would be reduced. Therefore, it can be deduced that *D_M_*=*D*+1=4, namely four-dimensional growth.

Mammals were representative of 3/4 dimensional evolution^4^. However, for protozoon and ectotherm, another key factor determining the metabolic rate was body surface area (*A*), which determined the respiration of protozoon and the maintenance of body temperature of ectotherm and therefore cannot be ignored. Based on similar methods, it was derived that the total metabolic dimension *D_B_*=4. The growth dimension *D_M_* remained unchanged, so^4^ *b*=4/4=1 for protozoon and ectotherm. More content can refer to Appendix 2.

Animals clearly showed a logic in metabolic macroevolution: after evolving the four-dimensional growth mode from the beginning of paramecium, how to control the systemic instability caused by body surface has become the biggest contradiction of the "spatial internalization" route in evolution. This therefore started the continuous evolution of the internal circulation system and body surface temperature control (including breathing). This contradiction had not been solved until the large reptiles, but caused greater weight burden and more dependence on the external environment (resources). From this perspective, the extinction of dinosaurs caused by major environmental events^35^ was accidental and inevitable. It was not until the evolution of the internal circulation system of thermostatic animals was completed, that the animals completely eliminated their dependence on body surface metabolism, and decreased again. A marker of the evolution to the top was the fractal of the internal circulation network^8^. In this sense, fractal growth was only a skill in evolution, rather than the general logic in macroevolution.

#### Comparison of b between empirical values and model derived value

It was speculated that *b* was among 3/4∼1 in animals, which was highly consistent with the existing actual observation data^1–4^. Similar to others, the *b* value derived from the priori model was not the maximum value of metabolic evolutionary selection, but the metabolic optimization eigenvalue that indicated the four-dimensional growth. Integrating the evolution logic of plants and animals, the final *b* value selected in macroevolution at the boundary level, both pointed to the top of secondary evolution.

### Fungi b = f (5) /5 = 5/5=1, D_B_=5, D_M_=5

#### Geometric derivation of the five-dimensional growth for fungi

Research on the metabolic scaling of fungi has been lacking for technological reasons. This study is also the first to mention the five-dimensional growth of fungi. Continuing the previous logic of dimension analysis, this paper makes the following analysis. More details can be seen in methods.

First, mycelium was a common choice of individual morphology evolution for fungi. Saprophytic mycelium could be simplified into a space growth mode in which individuals were fully filled in the three-dimensional nutrient space, as a living network^36^. Compared with animals, mycelium could be regarded as a monster whose whole body is a mouth. Therefore, the spatial generation modes of the growth dimension and metabolic dimension were completely consistent in the vegetative growth stage.

Second, the absorption function of fungi depended on the relative surface area of each cell linearly connected in space. They could maximize the relative surface area only through multicellular cooperation. At the same time, the evolutionary advantage of fungal cell size over bacteria was reflected (the width of fungal hyphae is approximately 5∼10 µm, which is several times to dozens of times larger than bacteria and actinomycetes), and therefore the absolute absorption surface area was also greatly increased. Obviously, this was a two-dimensional mode.

Finally, fungal hyphae could quickly form reticular branches through cell division and bifurcation links to fill the three-dimensional nutrient space. This growth mode was similar to the vascular growth behavior of animals and played the same three-dimensional effect^30^. Compared with "spatial internalization" of animals, it was actually to achieve barrier-free access to nutritional resources. Obviously, this was a three-dimensional mode.

In this way, higher fungi completed the optimization of spatial growth mode in two different metabolic key links (possession and absorption, and inner transportation function at the same time). Different from plants, these two links conformed to the relationship of multiplication. Therefore, there was

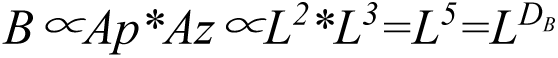

*Ap*, biological surface area that determines absorption efficiency, *Az*, biological surface area that determines acquisition efficiency. Therefore, *D_B_*=5. Mycelium is a structure of growth-metabolism fully integrated, therefore, *D_M_=D_B_*=5, and *b*=*D_B_/D_M_*=5/5=1.

#### Origin of five-dimensional growth in macroevolution

It was argued that there was a close relationship between one-dimensional evolution and five-dimensional growth mode, similar to the relationship between two-dimensional evolution and four-dimensional growth. More content can be found in the Appendix one A1.2. However, the origin and evolutionary path of fungi has been a complicated problem.

#### Comparison of b between empirical values and model derived value

Due to the limitations of actual measurement technology, the study of metabolic scaling for fungi has always been lacking, but there was no problem in the understanding of high metabolic efficiency for fungi. Due to the universality of mycelium structure in the fungal kingdom, we speculated that the variation range of *b* was narrower, almost near Of course, if multinucleated cells were taken into account, the actual value must be greater than 1.

Fungi at last evolved into "metabolic machine", compared with plants and animals. In summary, fungi, as the last one in the prior model, achieved the maximization of metabolic efficiency, and the final unification of structure and function evolution in metabolism.

## Synthesis and comparison of dimension application in macroevolution

The golden section is a natural law for the optimal utilization of space, and the Fibonacci function is an approaching function to the number of golden sections. Through the systematic derivation and verification of the prior model, it was argued that the macroevolution from prokaryotes to fungi was a highly order process of life emergence measured by the dimension application, and the realization trajectory of metabolic efficiency strictly followed the Fibonacci function.

Here, the five biological spatial growth optimization modes from prokaryotes to fungi in macroevolution were summarized and compared (Fig. 1 and Table 3). Strictly speaking, all of the metabolic dimension generation was derived from the surface area principle. The dimensional application of different life forms in macroevolution was derived from the geometric transformation and dimensional combination based on the surface area principle.

**Fig. 1.**
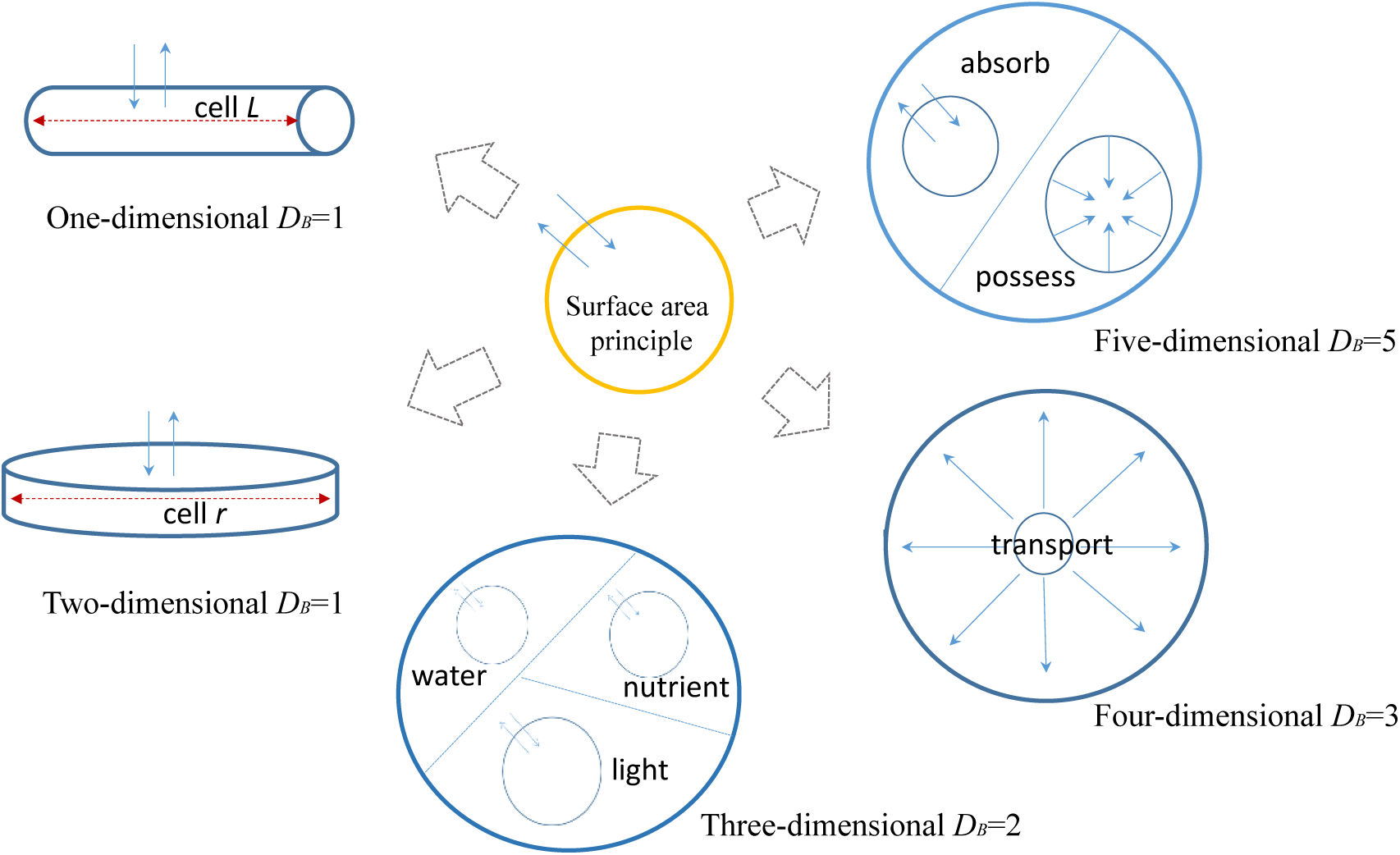
Five biological spatial growth optimization modes from prokaryotes to fungi in macroevolution. It referred to the pattern diagram of the metabolic scaling model^26^ and improved.

**Table 3.** Geometric operation rules of dimension application in macroevolution.

| Life forms | Variables |  |  |
| --- | --- | --- | --- |
| | Biology length $L$ | Biology area $A$ | Biology volume $V$ |
| Prokaryotes | $L \propto A \propto V \propto M \propto B$ | $A \propto L \propto V \propto M$ | $V \propto L \propto M$ |
| Protist | $L \propto A^{1/2} \propto V^{1/2} \propto M^{1/2} \propto B$ | $A \propto L^2 \propto V \propto M$ | $V \propto L^2 \propto M$ |
| Plants | $L \propto A^{1/2} \propto V^{1/3} \propto M^{1/3}$ | $A \propto L^2 \propto V^{2/3} \propto M^{2/3} \propto B$ | $V \propto L^3 \propto M$ |
| Animals | $L \propto A^{1/3} \propto V^{1/4} \propto M^{1/4}$ | $A \propto L^3 \propto V^{3/4} \propto M^{3/4} \propto B$ | $V \propto L^4 \propto M$ |
| Fungi | $L \propto A^{1/5} \propto V^{1/5} \propto M^{1/5}$ | $A \propto L^5 \propto V \propto M \propto B$ | $V \propto L^5 \propto M$ |

Prokaryotes and protists chose one-dimensional and two-dimensional growth routes to maximize the relative surface area of cells, respectively. The result was the geometric transformation from biological area *A* to biological length *L* for their metabolic dimension generation. Plants were a simple addition of multiple two-dimensional absorption processes of key resources (light, water, nutrition) without substantive dimensional combination or geometric transformation (more contents can refer to Method). Animals seemed to break through the surface area principle, and the metabolic efficiency was still determined by the cross-sectional area and total length of the pipe, realizing the geometric transformation from biological area *A* to physical space volume *V_D_*. Compared with plants, fungi were representatives of giving full play to the principle of surface area, realizing the true combination of dimensions and the perfect unity of growth dimension and metabolism dimension, but there was no substantive geometric transformation in this process.

## Verification conclusion

In macroevolution at the boundary level, the optimized growth mode in biological space from one-dimensional prokaryotes to five-dimensional fungi evolved in turn, accompanied by metabolic fluctuations and cycles from *b*=1 to *b*=1. Macroevolution was a selecting and optimizing process to utilize the physical space dimension and create biological dimensions, following the golden section law.

Evolution was generally regarded as a natural selection process for organisms to maximize metabolic efficiency in evolutionary ecology. However, it was shown that the principal line of macroevolution was not the realization of metabolism but the application of dimensions. The *b* values derived from the priori model have never been the maximum value of metabolism, but the characterization values of metabolic macroevolution were realized by dimension application and the indicator values of secondary evolution direction. Therefore, the original hypothesis that the *b* value was the indicator of maximum metabolic efficiency in macroevolution was not tenable. However, the variation in *b* indeed indicated some major evolutionary events from prokaryotes to fungi in macroevolution.

*b*=*D_B_/D_M_*=*f*(*n*)/*n* revealed the metabolic logic and dimensional logic of macroevolution, and therefore, *b* was given a clear physical definition and evolutionary connotation. It was argued that, there was such a biophysical model that metabolic scaling was unified with Fibonacci sequence in macroevolution, that is, the golden section model of macroevolution,

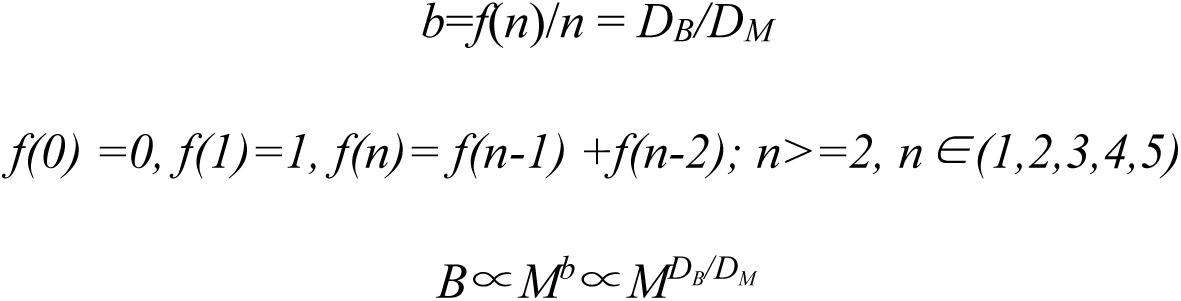

The mathematical form of macroevolution was a Fibonacci function with growth dimension as the independent variable and metabolism dimension as the variable, namely metabolic macroevolution function. Different from the general biological application of Fibonacci function, *n* was no longer a time unit but a dimension unit that determined the rhythm of macroevolution.

## Deduces

### Form and logic of metabolic evolution for cellular organisms

Fluctuation was the condition of life evolution, and convergence was the characteristic of life evolution. As shown in Fig. 2, metabolic evolution presented a general form of repeated fluctuations and gradual convergence. Macroevolution at the boundary level was open and closed, and the secondary evolution had ups and downs. Dimension reduction led to fluctuations, and dimension increase led to convergence. Evolution has never been a linear process, but a highly nonlinear process of layer-by-layer promotion (cell-individual-system), causal dependence (growth-metabolism), optimal selection (structure-function), and a highly ordered process (from one-dimensional to five dimensional) under the rule of golden section at the same time.

**Fig. 2.**
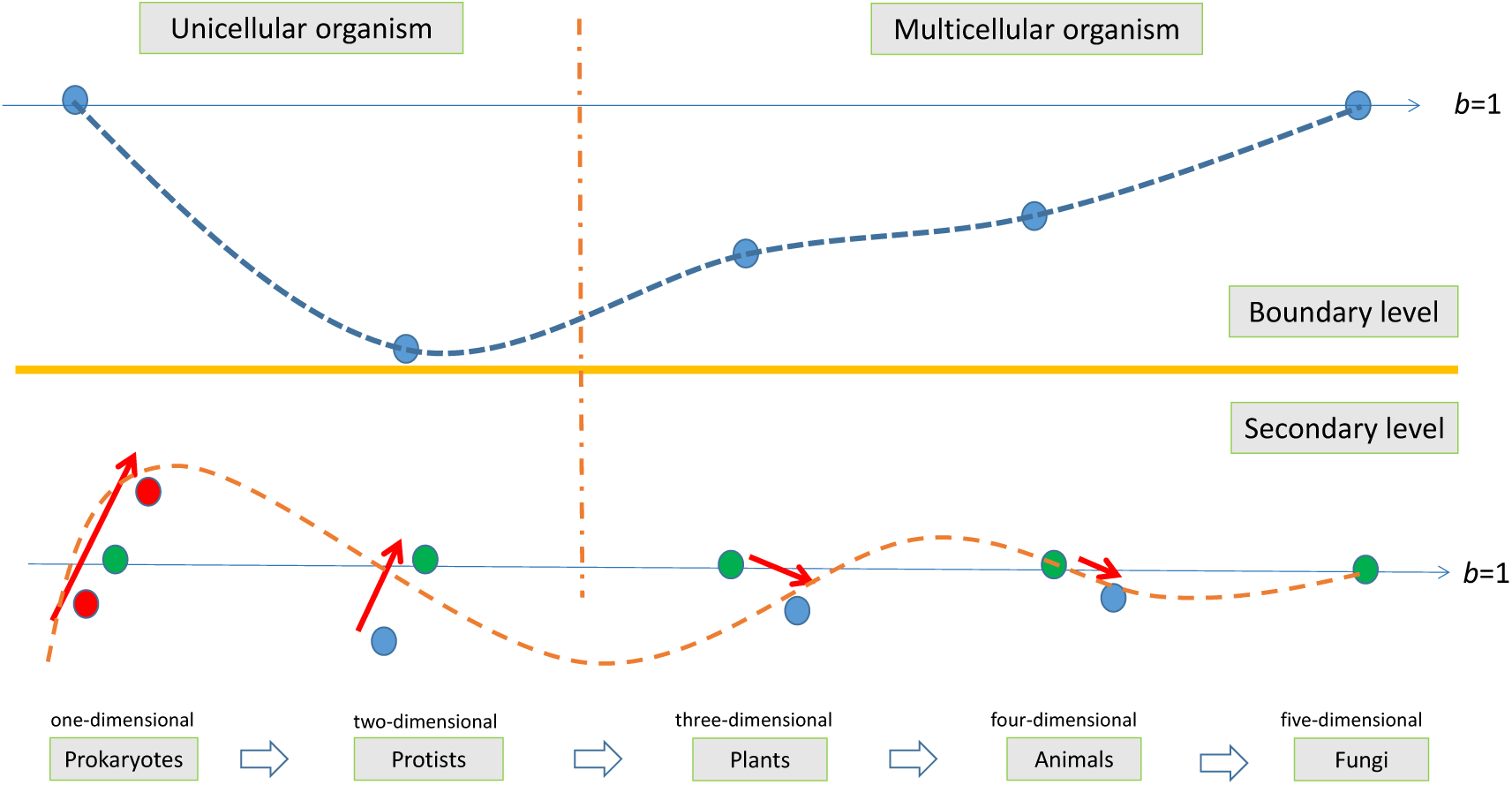
General form of metabolic evolution (i.e., b values) for cellular organisms in macroevolution. It is drawn based on the derivation results of dimension evolution in this paper and referring to the relevant research conclusions1–4. b, Characteristic value of metabolic scaling function (B ∝ M^b^). Boundary level, macroevolution at boundary level. Secondary level, macroevolution at levels lower than the boundary (such as phylum or class). The red arrows indicate the direction of b value evolution, and the length indicates their ranges. The points indicate the value of b, and the vertical position of the point indicates their relative sizes at the boundary level and at the secondary level, respectively. Two long solid lines with arrows indicate the position and trajectory of b=1. The two curves approximately fit the evolutionary trajectories of b at different levels.

The golden section was the spokesman of order and efficiency in nature. Life itself was an order emerging from disorder. For life, the pursuit of efficiency cannot escape the observance of order. In macroevolution, the molecular evolution of multinucleated cells played a key role in promoting fluctuations^2^, but the origin came from the one-dimensional growth in prokaryotes. Convergence started from the evolution of eukaryotic cells, but the two-dimensional growth of early protists was not unrelated to eukaryotic evolution. Metabolic homeostasis hypothesis^1^ and allometric scaling convergence hypothesis^2^ revealed the metabolic scaling laws across boundaries. However, the golden section law further became a rule to restrict the metabolic homeostasis law and allometry scaling convergence law. More content can be found in Appendix one (A1.1).

### Three-level metabolic evolution of cellular organisms

The unification of metabolic scaling and *Fibonacci* sequence in macroevolution provided a full-new perspective for us to rediscover the evolutionary tree of life from views of metabolic evolution and dimension evolution.

As shown in Fig. 3, there were three levels of metabolic macroevolution. The first evolution was from prokaryotes to protists, marked by the emergence of eukaryotes, which could be called the trade-off stage of dimension application and metabolic achievement. The second evolution was the differentiation of plants, animals and fungi from unicellular eukaryotes, marked by the coevolution of the individual structure and function of higher plants, higher animals and higher fungi, which could be called the synergistic stage of dimension application and metabolic achievement. The third evolution was marked by the fact that fungi were at the top of eukaryotic metabolic efficiency and completed the evolution of ecosystems, which could be called the upgrading stage of system evolution and metabolic cycle.

**Fig. 3.**
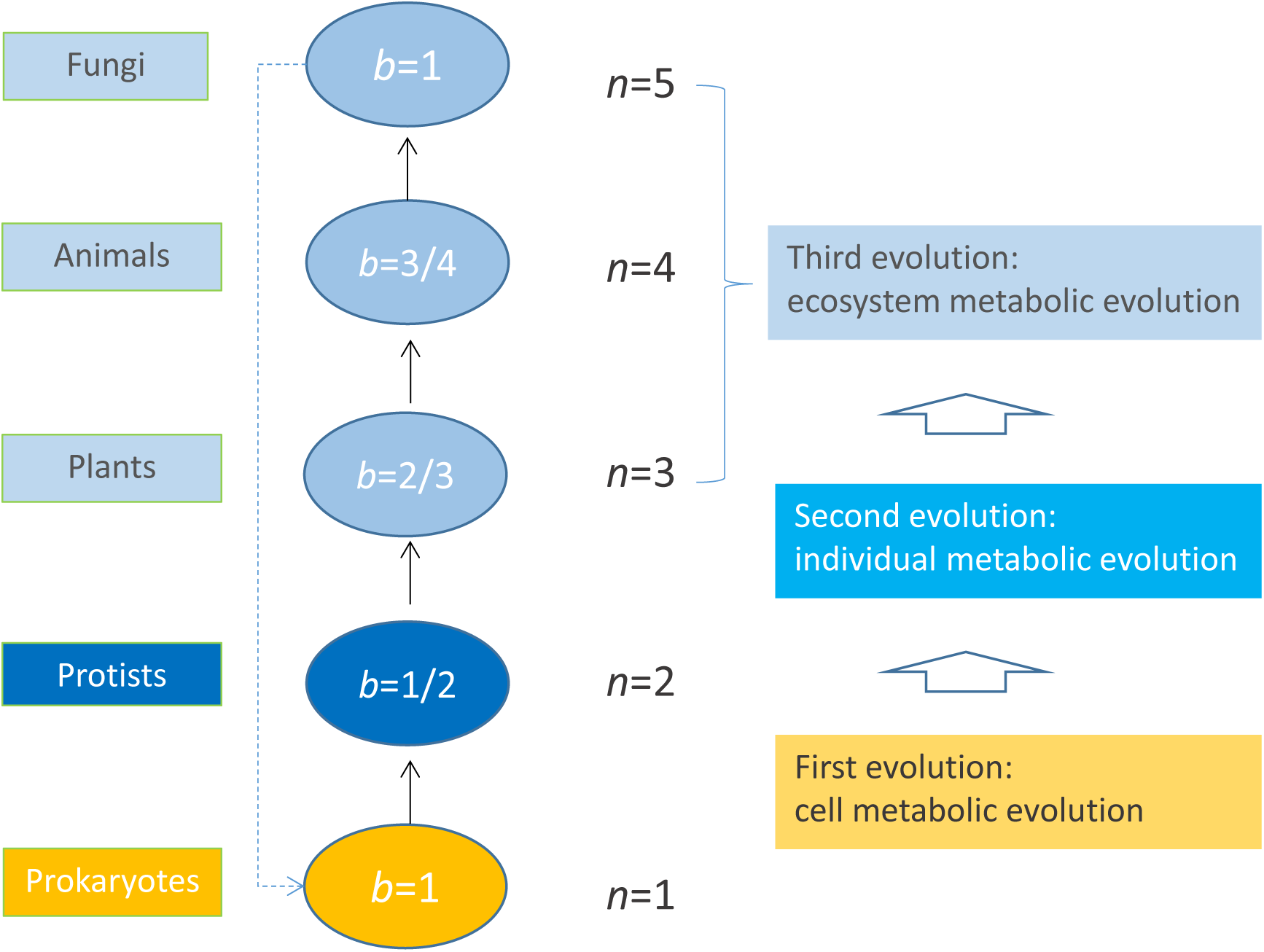
Panorama of three-level metabolic evolution of cellular organisms. **b**, Characteristic value of the metabolic scaling function *(B∝Mb). **n***, independent variable in Fibonacci function. More contents can refer to appendix one (A1.2).

### Evolutionary logic and clues in the early stage of macroevolution

The golden section model of macroevolution revealed another key evolutionary mechanism, namely dimensional evolution. As shown in Figure 4, based on the research of macroevolution^20, 37, 38^, molecular evolution^25, 27, 34, 39^, and the idea of dimensional evolution proposed in this paper, the logic of early life evolution was re-combed, and three paths of early evolution of higher life, as well as the key evolutionary events based on the interaction between dimensional evolution and molecular evolution, were proposed.

**Fig. 4.**
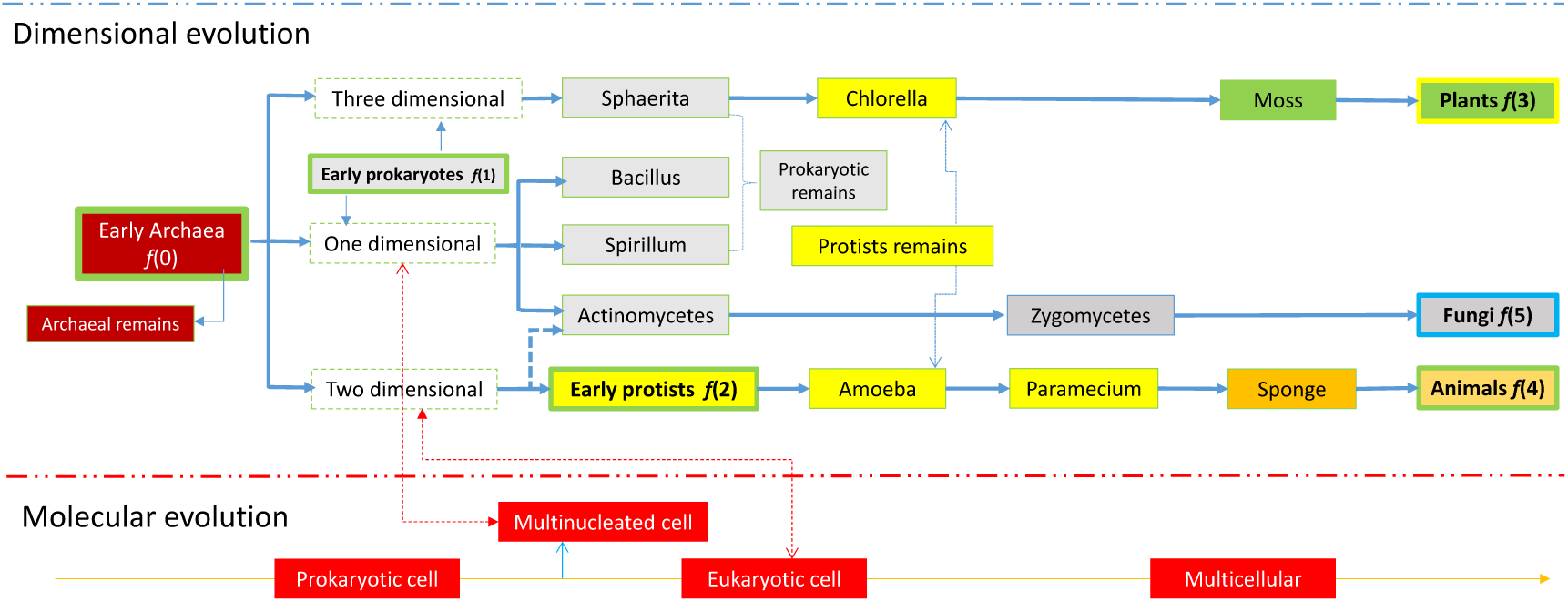
Overall trajectory and path differentiation in the early evolution stage. More content can be found in Appendix one (A1.2).

Here, early archaea was taken as the common ancestor of prokaryotes and protists. Taking dimensional evolution as the main line, early archaea differentiated into three routes, namely, one-dimensional, two-dimensional and three-dimensional conservative evolution. Ones taking the one-dimensional and three-dimensional routes developed into prokaryotes, in which the three-dimensional ones might evolve through independent molecular evolution of eukaryotic to plants. The two-dimensional ones entered the fast track of eukaryotic molecular evolution and first completed the evolution of eukaryotic cells, which in turn became the starting point of the evolutionary path of animals. The molecular evolution of multinucleated cells induced by one-dimensional evolution became the starting point of the fungal evolution route.

At the same time of dimensional differentiation, molecular evolution was the auxiliary line, multinucleate cell evolution was intertwined with one-dimensional evolution, and eukaryotic cell evolution was intertwined with two-dimensional evolution, which constituted the key events and nodes in the early evolution stage.

### Extended reasoning

It also extended and discussed the significance and essence of the golden section to life and evolution, the form and logic of dimensional evolution in evolution, the unification of the metabolic scaling model, the geometric basis of macroevolution analysis, and the *Fibonacci* code of macroevolution including viruses and the nervous system. More content can be found in Appendix one.

### Summary

In any case, the hypothesis indeed provided a powerful explanation for the empirical research results both in metabolic scaling and evolution studies, and provided a full-new logic and ways to integrate existing experience and theoretical research.

1. Through the integration of macroevolution and golden section law, it was found that the macroevolution of the cell life system was a highly organized process from one-dimensional to five-dimensional evolution. This has broken the long-standing understanding of macroevolution: a highly nonlinear, complex interactive system far from equity. Thus, it also raised a basic problem, what was the basic yardstick to measure macroevolution? This actually reflected the essence of the macroevolution process. It was argued that the golden section gave a clear rhythm and connotation to macroevolution, not time, but the biological space dimension related to the metabolic scaling rule.
2. Through the integration of metabolic scaling and Fibonacci series, it was found that the *b* value was actually the result of selecting optimization between the metabolic dimension and growth dimension, corresponding to different life forms in macroevolution. At the same time, the variation in the *b* value indicated major evolutionary events from prokaryotes to fungi in macroevolution, and defined the direction of secondary evolution. Thus, it also revealed another important logic of macroevolution, that is, metabolic evolution.
3. Compared with the current highly diverse and complicated molecular evolution evidence and fossil evidence, both dimensional evolution and metabolic evolution revealed the potential clear and orderly logic in macroevolution. With dimensional evolution as the main line and molecular evolution as the auxiliary line, it was argued that the golden section model of macroevolution provided a possible path for the unification of macroevolution and microevolution.

## Acknowledgements

We thank many contributors to the metabolic scaling database and geometric model. We also thank *Prof* Y.X. WANG, *Prof* X. CHENG, *Prof* N.P. SONG for their supports.

## Author contributions

X.Y developed the design of the study, constructed the model and wrote the text. L.W improved the geometric analysis method.

## Method

### Concept and method of macroevolutionary geometric analysis

#### Optimized growth mode in biological space

"Optimized growth mode in biological space" refers to the spatial generation of biological dimensions, including the spatial geometric growth of the individual itself and that of "functional components" that determine individual metabolic efficiency. This is the concept of the use of spatial dimensions by biological growth and metabolic activities, including the growth dimension *D_M_* and metabolic dimension *D_B_*.

#### Geometric transformation

In the evolution of life, the spatial growth mode of "functional components" of organism metabolism may switch in one-dimensional, two-dimensional and three-dimensional physical space, so that a transformation of geometric operation rules must be done, since the surface area principle that determines the metabolic rate no longer strictly follows the Euclidean space geometric algorithm.

#### Dimension combination

When multiple different metabolic links limit the realization of metabolic efficiency at the same time, individuals will evolve different spatial growth patterns and corresponding dimension applications for different metabolic links. At this time, the total metabolic dimension of an individual is the addition of the dimensions of each link (expressed in geometric operations, it is the multiplication of biological areas), such as fungi. However, the absorption of different types of restricted resources is only a single metabolic link. The total metabolic dimension is the mean of the dimensions of spatial growth patterns of different resource absorption functional components (expressed in geometric operations, it is the addition of biological areas), such as plants.

#### General method

Referring to the definition of dimension concepts related to biological growth (individual size) and metabolism^5^, it is set *b*=*D_B_*/*D_M_*. *D_B_*, metabolic dimension, *D_M_*, growth dimension. Due to the limitation of the physical dimension, *M* is always proportional to biological volume *V*, and there is *V∝L^D_M_^*. When *B* is proportional to biological area *A*, there is *A ∝ L^D_B_^*. *L*, biological length. Some restrictions in geometric operation are mentioned in the following:

1. Limited by the structure and function of cellular organisms, the physical dimension of individuals *D*=3, but the biological dimension of individuals (including *D_B_* and *D_M_*) is determined by the spatial growth pattern. For cellular organisms (without considering the influence of nonspatial factors on metabolism) in macroevolution, *D_B_* is always a part of *D_M_*, less than or equal to *D_M_*, but the difference between them will not be greater than 1 in general, according to the metabolic optimization principle and the law of mass conservation^30^.
2. In metabolic macroevolution, the nutritional environment (supply mode) and different metabolic links may become the key to determining the geometric operation of the metabolic dimension and growth dimension. For unicellular organisms, both the coordination of multiple metabolic organs and the relative surface area of cells may become key limiting factors. Therefore, *B* is not always isometric to *A*.

#### Biological length, biological area and biological volume

The connotations of biological length (*L*), biological area (*A*) and biological volume (*V*) were expanded and summarized (table. 4), based on the summary of biological dimensions by West et al.^5^, and combined with the experience of biological geometric analysis in the macroevolution.

**Table 4.** Biological length (*L*), biological area (*A*) and biological volume (*V*) in metabolic macroevolution geometric analysis.

| Variable | Life forms in macroevolution |  |  |  |
| --- | --- | --- | --- | --- |
|  | Unicellular organism | Plants | Animals | Fungi |
| $L$ | cell radius or thickness; long axis of cell; distance between organelles; etc. | individual height; leaf canopy diameter; leaf canopy height; length parameters of absorbing roots; etc. | length of blood vessels, the distance from the heart to the body surface, and the length and thickness of the body. etc. | length of mycelium; cell radius or thickness; long axis of cell; |
| $A$ | Cell surface area; etc. | leaf canopy area; absorbing roots system; etc. | Body surface area; Vascular cross-sectional area; etc. | Cell surface area; etc. |
| $V$ | Cell volume; etc. | individual volume; roots system; etc. | Body size; Vascular system; etc. | Cell volume; mycelium system; etc. |

Among all the parameters, biological length is the most critical and easily misused basic geometric parameter. West et al.^5^ defined connotations of biological length focused on the metabolic network. From the perspective of more general metabolic constraints, for unicellular organisms, it should first include the cell radius or thickness, the length of long axis of cell, the distance between multi-organelles, etc. For plants, biological length parameters that restrict metabolism mainly include individual height (which determines the distance between leaves and roots), leaf canopy diameter, leaf canopy height, and length characteristic parameters related to the dense distribution layer of absorbing roots. For animals, the most basic is the length characteristic parameters of the internal circulation network, such as the length of blood vessels (digestive tract), the distance from the heart to the body surface, and the length and thickness of the body. Fungi are complex and simple. The most basic is the length of mycelium (which determines the distance from the nutrient source). In addition, the length parameters at the cell level (which is determined by the special metabolic structure of mycelium), such as the length of the long axis of the cell, the radius or thickness of the short axis of the cell, etc.

Area and volume are defined by length. However, it should be emphasized that the geometric relation between leaf canopy and plant individuals will change due to the different morphological evolution and growth stages of plants, since plant is a kind of modular organism with high growth elasticity. Fungi is another special case. Both of them make the macroevolutionary geometry operation surpass the general Euclidean geometry and fractal geometry operation rules^5^.

### General geometric model of dimension analysis in macroevolution

#### Unicellular organisms

Let cell volume be *V*, cell surface area be *A*, cell radius be *r*, and biological length be *L*.

#### Linear cells (one-dimensional growth cells)

Prokaryotes with one-dimensional growth can be approximately regarded as a two sharpened pencil whose bottom area can be ignored. In evolution there is a minimum internal radius of cell *r* due to the limits of cell structure and function. *r* is approximately constant in the elongating growth of the cell, and *r* is very small compared with cell length *L*. Therefore, from *A*=2π*rL* and *V*=π*r*^2^*L, r* is constant, it can be concluded that *V∝A∝L*. From *A*∝*B*∝ *L^D^*^B^, *V*∝*M*∝*L^D^*^M^, there is *D_B_*=1, *D_M_*=1, and *b*=*D_B_*/*D_M_*=1/1=1. *A*, biological area. *V*, biological volume. One-dimensional growth can be considered a specific case of the metabolic boundary restriction hypothesis^13^ in geometric derivation.

When the cell grows fully and radially in a plane with a radius of *L* (such as actinomycetes), there is *V*∝*l*^2^, *A*∝*l*^2^, *L*∝*l*, so *D*_M_=2, *D*_B_=2, *b*=*D_B_/D_M_*=2/2=1.

#### Cake shaped cells (two-dimensional growth cells)

Protists with two-dimensional growth can be approximately regarded as a squashed rugby whose side area can be ignored. In evolution there is a minimum internal thickness *h* due to the limits of cell structure and function. *h* is approximately constant in the horizontal extension of the cell, and *h* is very small compared with cell radius *r*. From *A*=π*r*^2^, *V*=π*r*^2^*h, h* is constant, it can be concluded that, *V∝A∝r*^2^*∝L*^2^, there is *V∝L*^2^*∝L^D_M_^*, *D_M_*=2. However, for protists with two-dimensional growth in some critical state, the metabolic efficiency is determined by the biological length *L* (i.e., *r*), rather than *A*, that is *L∝L^D_B_^*, therefore, *D_B_*=1, *b* = D*_B_*/D*_M_* = 1/2.

When metabolic efficiency is limited by both cell surface area and internal distance, that is, when the cell morphology is between three-dimensional and one-dimensional, there is no simple geometric operation model, but the range of *b* value^13^ is generally in 2/3∼1.

#### Globular cells

There are *V∝r*^3^*, A∝r*^2^*, L∝r*. When metabolic efficiency is determined by cell surface area, *A∝r*^2^*∝L*^2^*∝B*. Under the condition of uniform cell density, *V∝r*^3^*∝L*^3^*∝M*. From *V∝L^D_M_^, A∝L^D_B_^*, *b*=*D_B_*/*D_M_*, *D_M_*=3, *D_B_*=2, and *b*=2/3.

When metabolic efficiency is determined by cell volume^13^ (for example of *Chlorella*, chloroplast particles that determine metabolic efficiency are filled in three-dimensional cells), there is *V∝M∝r*^3^*∝L*^3^*∝A∝B*, so *D_M_*=3, *D_B_*=3, *b*=3/3=1.

#### Animals

The least carrier theory^30^ reveals the general logic and basic spatial growth mode of dimension evolution in macroevolution for animals. Based on this, this paper constructs a unified geometric model of animal dimension application and metabolism realization (please see Appendix 2 for the specific derivation process).

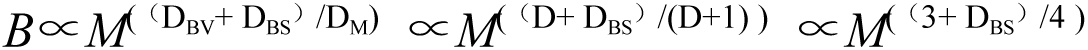

Among them, *D_BV_* is the basic metabolic dimension (i.e. the spatial dimension of the internal circulation system), *D_BS_* is the surface metabolic dimension, *D* is the physical spatial dimension occupied by individuals, which is equal to 3, and *D_BV_=D*. *D_BS_*=1 for low and variable temperature animals, and *D_BS_*=0 for constant temperature animals with fully developed internal circulation.

#### Plants (tall and mature trees)

This paper constructs a basic geometric model of plant metabolic macroevolution by integrating the minimum vector theory^30^, the plant extension of the WBE model^40^, and the classical tree mechanics model^28, 29^.

Let *Q* be the xylem sap flow rate (v/d), *Q_M_* be the total xylem sap volume in a day, *D* be the trunk diameter, and *H* be the height of the tree. there is^40^ *Q* ∝ *D*^2^. The total xylem sap volume measured in a day is equivalent to the xylem sap flow rate. Therefore, *Q_M_∝Q∝D*^2^ exists. In terms of macroevolution, according to the principle of metabolic optimization, with reference to the least carrier theory and the law of mass conservation^30^, the higher the transmission efficiency of the network, the less the capacity of the carrier is required (the total liquid volume of xylem and animal blood play the same carrier role), *M*∝*Q_M_*∝*D*^2^ can also be deduced (*Q_M_*∝*M*^25/24^ was also deduced^40^), rather than^9^ *B* ∝ *Q* ∝ *D*^2^. From *H* ∝ *D*^2/3^, *M* ∝ *H*^3^ can be obtained. Therefore, tall trees at the top of plant evolution are typical representatives of three-dimensional growth patterns, following the traditional Euclidean space geometric algorithm. From *M*∝V∝*L*^3^, *B*∝A∝*L*^2^, *B*∝*M*^2/3^ can be directly obtained. Please refer to Appendix 3 for the specific derivation and analysis process.

#### Multi-Resource-constrained geometric derivation of the three-dimensional growth

The efficient use of light for plants was a unique principle in evolution, and the mode of resource supply therefore became the basis of space growth selection for plants^17^. The -3/2 self-thinning model^15^ revealed the scaling between a two-dimensional surface (leaf canopy) and the three-dimensional individual, following energy equivalence^31^.

Water and nutrients may also become the limiting factors of metabolism. Most of the available nutrients were concentrated nearly in the shallow soil layer, which can be approximately regarded as a plane relative to the whole volume of tree individuals. Water, originating from precipitation, was also a two-dimensional input mode similar to light. In this way, the supplies of various restricted resources for plants could be considered nearly two-dimensional.

The spatial growth mode of tall and mature trees can be imagined as a bundle of tightly bound pipes connected up and down, and the functional components for the absorption of light, water and nutrients were located at both ends of these tubes. Limited by the space positional relationship, the space growth mode of the leaf canopy and the absorbing root system, both could be nearly considered two-dimensional compared with the growth of tree individuals. Moreover, the resource supply pattern and the metabolism-related growth mode therefore became coincident in the space dimension, which was a reasonable choice in evolution for plants.

Although the original single light limitation was changed to light, water and nutrients, the two-dimensional growth of absorption components remained unchanged, and the metabolic link that determined the metabolic rate remained unchanged(i.e., absorption). Therefore, there was only a simple additive relationship in mathematics in the geometric operation of the metabolic dimension as follows:

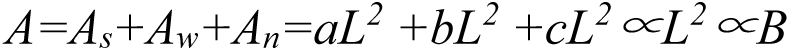

*A*, the total biological area, *A_s_*, the biological area of light absorption, *A_w_*, the biological area of water absorption, *A_n_*, the biological area of nutrient absorption, *a*, *b* and *C* are constants. *L*, biological length.

#### Fungi

For the vegetative growth stage of fungal mycelium, there are two key links restricting metabolic efficiency at the same time: possession (A_Z_)and then absorption (A_P_). According to the principle of dimension combination, *B*∝*Ap*\**Az*.

Let the mycelium volume be *V* (the space occupied), the surface area be *A* (the total surface area of multicellular bodies), and the characteristic length be *L* (the average distance from the starting growth point of spores to the farthest end). The dimension of the physical space where the nutrient is located is *D*, which is equal to 3. The possession is determined by the volume of the mycelium, that is, *Az*∝*V*. When the mycelium is nearly filled in the physical space where the nutrient is located, there is *V* ∝*L^D^*, so *Az∝V∝L^D^∝L*^3^.

Similarly, the absorption is determined by the total surface area of mycelium, that is, *Ap*∝*A*. The surface area of each cell *A_C_*∝*r^2^*, *r*∝*L, r*cell radius, and the number of cells is *N*. For the space growth mode of mycelium linked by many single cells, there is *A*=*A*_c1_+*A*_c2_+……+*A*_cN_=*ar*^2^+ *br*^2^+……+ *nr*^2^∝*r*^2^∝*L*^2^, where *a, b, x* are constants.

Therefore, *B*∝Ap*Az∝*L*^2^\**L*^3^=*L*^5^, D_B_=5. Because the spatial pattern of individual mycelium growth and metabolic component growth was completely integrated, *D_M_=D_B_*=5, and *b=D_B_/D_M_*=1.

## Appendix one. Extended reasoning and supplementary discussion

### A1.1 Essence of golden section in macroevolution

#### Golden section and macroevolution

Mathematically, golden section referred to dividing the whole into two parts. The ratio of the larger part to the whole part was equal to the ratio of the smaller part to the larger part, and its ratio is about 0.618. Golden section just is a dichotomy of nature. Figuratively speaking, golden section divided space into two parts in metabolic macroevolution, one for growth and the other for metabolism. In macroevolution, the first five knives of golden section cut out five life forms in turn, leaving a clear trace, that was, the dimensional evolution from one to five dimensional space growth modes.

The method to calculate the golden section number (0.618) is to calculate the ratio of two adjacent numbers from the second place of Fibonacci sequence 1, 1, 2, 3, 5, 8, 13, 21,…, that is, the approximate value of 2/3,3/5,5/8,8/13,13/21. In fact, another wonderful function of golden section is the ratio of independent variables to variables, just as did in this article.

The essence of golden section to macroevolution, from the analysis results of this paper, was that life used the physical space dimensions and independently created biological dimensions to realize metabolic optimization. All the questions were not why macroevolution chose golden section, but why golden section chose macroevolution. To answer this question, there was a more ultimate question, where did golden section come from? From the cantilever structure of galaxy, to the arrangement of petals of a flower, and even the combination of some macromolecules, all followed golden section. Why? If this question can’t be answered, let’s taken it as an axiom of the universe firstly.

However, combined with the analysis of relevant research literature, we found that the role of golden section in metabolic macroevolution ultimately seemed to point to the mapping of the basic physical and chemical mechanisms of metabolic activities at different levels. For examples of, multinucleated cells (genetic material and energy expression) and one-dimensional^2^, material diffusion (chemical penetration theory) and two-dimensional^19^, mechanical restriction (biomechanics) and three-dimensional^28, 29^, fractal geometry (Fluid Mechanics) and four-dimensional^8^, as well as the surface area principle of metabolism (interface theory) and fungal five dimensionality^26^, quantum metabolism (quantum physical mechanism of energy metabolism) and differentiation of 2/3 and 3/4 rule^34^, etc.

Are there inevitable connections between these physical and chemical mechanisms that played a crucial role in microevolution and the application of dimensions dominated by golden section in macroevolution? Thinking about this problem may contribute to the understanding of the deep-seated connotation and potential sources of golden section.

#### Fibonacci function and macroevolution

##### Top five positions in Fibonacci series and five realm system of cellular life

Five realm life system was the mainstream classification method at present, which had a long history. From the philosophical and physical aspects of the origin of life, order was produced in disorder, and life was conceived in fluctuation, which was the definition of the physical essence of life system by non-equilibrium thermodynamics. The fluctuation-convergence properties of Fibonacci function were mainly concentrated in the top five positions (Fig. 5). In the Fibonacci function of life evolution, *n* was no longer a simple number, and *f* (n) was no longer a simple function operation result. Was this a coincidence? So our classification and ranking of five realm life system was also a coincidence? The earliest basis of the classification just was the biological differences of morphology and structures, which nicely proved that golden section was in the same line with macroevolution, since there was a close relationship between morphological evolution and dimensional evolution.

##### Approach properties to golden section point of Fibonacci function and the metabolic fluctuation and convergence in macroevolution

Comparing between Fig.2 and Fig. 5, we can find the similarity between Fibonacci function and metabolic evolution: first fluctuation and then Convergence. In macroevolution, molecular evolution of multinucleated cell played a key role in promoting fluctuations^2^, but the origin came from the one-dimensional selection in prokaryote. Convergence started from the evolution of eukaryotic cells, and the space growth mode of early protist was not unrelated to eukaryotic evolution. In short, the final result was that the trajectory of metabolic macroevolution^1–4^ strictly followed the approach trajectory to golden section point of *Fibonacci* function.

**Fig. 5.**
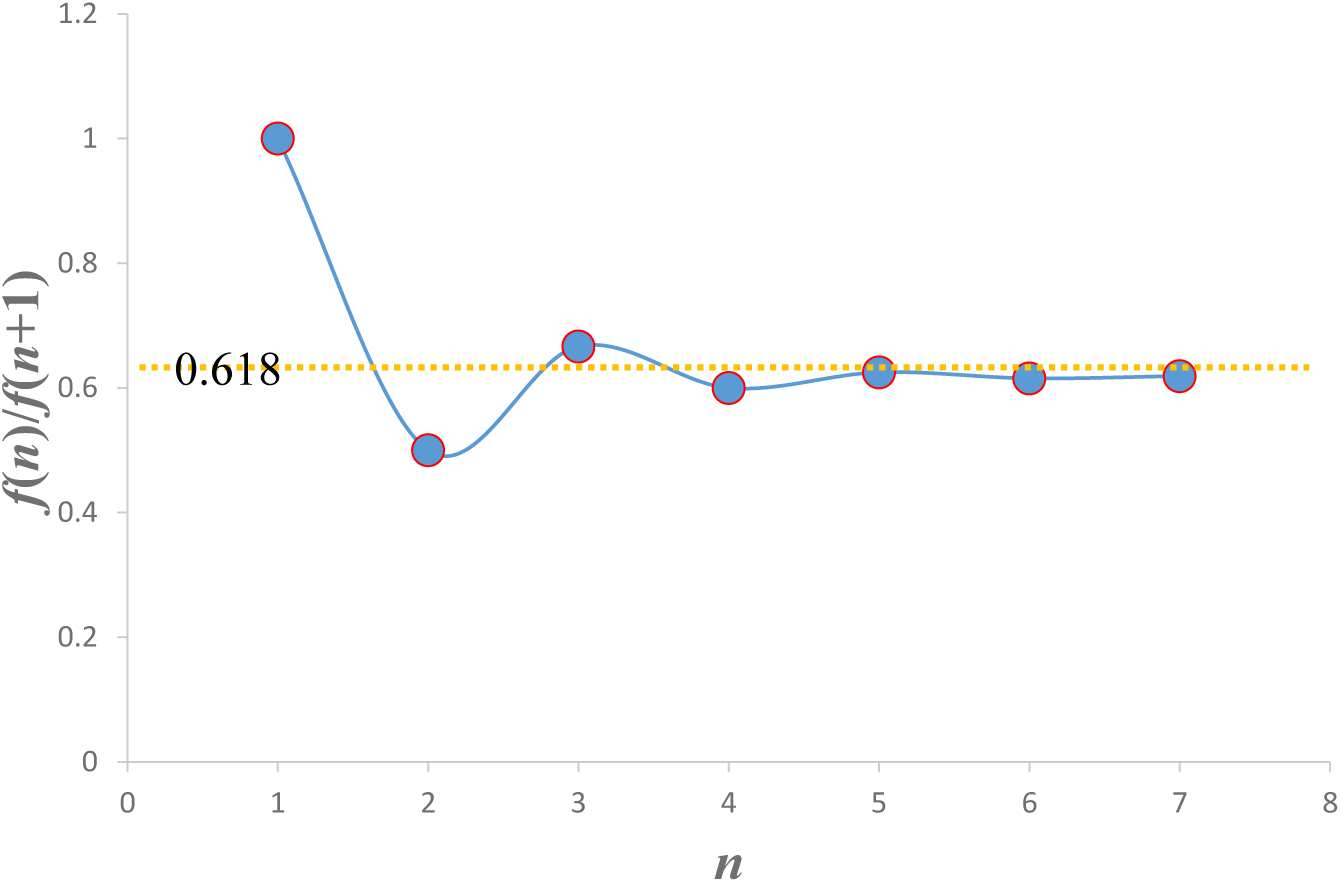
Golden section approach trajectory in Fibonacci function.

##### Inheritance operation and life evolution tree

An interesting problem was that according to *f* (*n*) =*f* (*n*-1) +*f* (*n*-2), the metabolic dimension of the latter life form should be the sum of the first two. However, it might reflect a logic of macroevolution in possible. The evolutionary isolation of life forms appeared at a rhythm that conformed to the progressive principle of Fibonacci function (exponential function based on *e*), which was the most efficient emergence of life forms? Maybe it can be started with the essence of natural logarithm *e* and make some further speculation. Of course, another possible speculation was that there were dimensional evolutionary relationship among some life forms with parasitic relationships (for example, most of early fungi must be parasitic). In short, all of modern higher life forms originated from unicellular eukaryotic. Understanding what happened at that stage may be the key to cracking the golden section password of macroevolution.

##### f (0), f (1), f (2) and the early life forms in evolution

The first two values in Fibonacci sequence were directly assigned, namely 0 and 1, but these two assignments were not optional, or which can be large or small. In macroevolution, prokaryote (*n*=1) was directly assigned a value of 1. *f* (0) may be archaea (*n*=0), but it was more likely to be the ancestor of archaea remains. This potential life form should not have the ability of dimensional evolution (*f* (0) =0). The evidences of molecular evolution showed that the genetic relationship between archaea, prokaryotes and protist were widely intertwined^25, 27, 39^. It was looked forward to further study of Archaea to find more evidence.

Another interesting question was that the early unicellular life stage represented by prokaryote (*f* (1) =1) and protist (*f* (2) =1) just was a gestation period of advanced life forms, which was very appropriate to the connotation of “rabbit sequence”, as the original prototype of Fibonacci sequence. It was an interesting association.

### A1.2 Significance of dimensional evolution in macroevolution

#### Evolutionary dichotomy: dimensional and molecular evolution

Dichotomy could be found everywhere in evolution and evolutionary analysis, for examples of, the fluctuation and convergence of evolution, metabolic evolution and growth (size) evolution, dimensional evolution and molecular evolution, growth dimension and metabolic dimension, and so on, even including microevolution and macroevolution, internal and external causes of evolution, etc. This reflected a basic model of evolution: evolution always repeatedly selected and then optimized in two contradictory aspects, just as the cell optimal growth theory^11^ revealing the selection optimization between cell number growth and size growth.

Golden section to metabolic macroevolution was actually a dichotomy of growth and metabolism. The random drift theory in molecular evolution was still a dichotomy (or two-sided bet, no tendency of advantage and disadvantage in molecular evolution). Dichotomy to evolution was the methodology of evolution itself. Similarly, it was also the methodology of evolution analysis.

#### Was the yardstick to measure macroevolution time or dimension?

Due to the long-term cognitive solidification brought by the complex molecular evidences and fossil evidences, macroevolution has been considered as a highly nonlinear, complex interactive system far from equilibrium, but at the same time it was also dominated by a simple natural law. However, this trajectory was not measured by time, but dimension. On the scale of billions of years, time was not an ideal yardstick to measure macroevolution, and it was indeed the case^41, 42^.

The latest research found that^37^, on a time scale between 40 and 500 *Myr*, biodiversity was increasingly insensitive to large fluctuations in climate status. This indicated that macroevolution always had a set of endogenous mechanisms independent of the external environment events. It was argued that this endogenous mechanism was determined together by the bottom drive of molecular evolution in microevolution, and the upper constraint of dimensional evolution in macroevolution.

This still was an evolutionary dichotomy.

#### Application of dimensional evolution to some issues in macroevolution

##### Evolutionary relationship among archaea, prokaryote, and protest

Origin of eukaryotic cells has been one of the most controversial problems in modern biology. Spang et al.^25^ provided support for the emergence of eukaryotic cells from archaea. It described a new candidate archaea, which genome encoded a series of extended eukaryotic characteristic proteins, suggesting complex membrane remodeling ability. Before it was speculated the potential role of two-dimensional evolution in eukaryotic cell evolution, such as cell membrane folding, and then speculated that there was a two-dimensional route in the early evolutionary stage, which eventually became the fast track of eukaryotic cell evolution (Fig.4). The evidence of molecular evolution^25^ basically affirmed our speculation.

Molecular evolution^25^ was intertwined with the dimensional evolution, and the great oxidation event finally led to the emergence of eukaryotes marked by sudden increase in size^20^, which might be a complete evolutionary logic of the origin of eukaryotic cells. Thus, the evolutionary relationship between archaea, prokaryote and protist based on dimensional evolution, should be tenable in evolutionary logic and key evidence (Fig.4).

##### Origin and evolutionary routes of plants, animals and fungi

Hedges et al.^27^ studied the molecular time scale of eukaryotic evolution and the rise of complex multicellular life. The results showed that the relationship between animals and fungi was closer than that with plants, the relationship between red algae and plants was closer than that with animals or fungi, and the relationship between flagellates and animals was closer than that with fungi or plants, Alveoli were the basis of fungi. This was basically consistent with our prediction of the origin and early evolutionary route of plants and animals based on dimensional evolution (Fig.4).

The origin and evolution of fungi has always been unclear and complex^43, 44^. Leander and Keeling^45^ found that morphological and molecular evidences showed that alveoli had a common ancestor, and together form one of the largest and most important eukaryotic microbial assemblages recognized today. But who was this ancestor? Under the logic of dimensional evolution, it was speculated that actinomycetes (or similar biology) may be this ancestor, since there were so much similarities between fungi^36, 44^ and actinomycetes, but it was still needed to clarify the molecular phylogenetic relationship between actinomycetes and alveoli.

From the initial one-dimensional evolution to mycelial route, it was an important and interesting evolutionary branch. It was speculated that Actinomycetes was an origin of an intermediate route integrating one-dimensional and two-dimensional routes (Fig.4), just as the intermediate transition evolution in alveoli^45^. Inspired by the research findings on red algae and red blood cells^39^ and the Cross-species gene transfer in fungi^44^, it was speculated that there was a mixed evolutionary route for early actinomycetes (mainly parasitic types) and early ancestor of animal, until the early fungi were isolated from this route and evolve independently, based on the earlier study results^27, 45^.

##### Ecosystem evolution in macroevolution

Macroevolution needed to answer a question, what was the co-evolutionary logic of plants, animals and fungi? Why did plants and animals evolve from simple (structure) but efficient (metabolism), to complex but relative inefficient? Why had fungi always followed the evolutionary strategy of maximizing metabolic efficiency, but their basic cell structure had not undergone significant differentiation as same as plants and animals?

To answer those questions, some inspiration might be got from the latest progress of macroevolution theory based on internal or external driving in recent years. First of all, biological drivers (internal driving theory) mainly involved the peak of taxonomic expansion, while non-biological drivers were mainly applicable to taxonomic extinction^46^, especially by driving the synchronous extinction of lineages originated from the more lasting and complex interaction between biological and environmental factors to affect evolution, rather than directly affected the formation rate of new species^35^. Moreover, the components determined by internal factors would also be more important on a larger evolutionary time scale^37^.

That was to say, macroevolution had a set of endogenous mechanisms independent of the external environment. First of all, in the cross-boundary evolution on the scale of billions of years, the interaction and coevolution among plants, animals and fungi cannot be ignored as one of the inner driving force of macroevolution. Secondly, there might be a logic of systematic evolution, so that the evolution of ecosystems was enough to buffer or overcome the synchronous extinction of pedigrees driven by external factors^35^. It was also believed that from cell evolution, individual evolution to ecosystem evolution, there might be a hierarchical evolution of metabolic homeostasis^1^ in macroevolution. Ecosystem evolution, why not?

### A1.3 Unification of metabolic scaling theory

#### Unified framework of metabolic scaling theory^26^ and its application in this paper

As shown in Figure 6, in this review, Glazier^26^ summarized and refined four theoretical paradigms of metabolic scaling, and proposed a unified framework of metabolic scaling theory (CMT). CMT recognized the openness, dynamics and complex hierarchy and interactive organization of biological systems, the importance of biological regulation of resource supply and demand.

**Figure 6.**
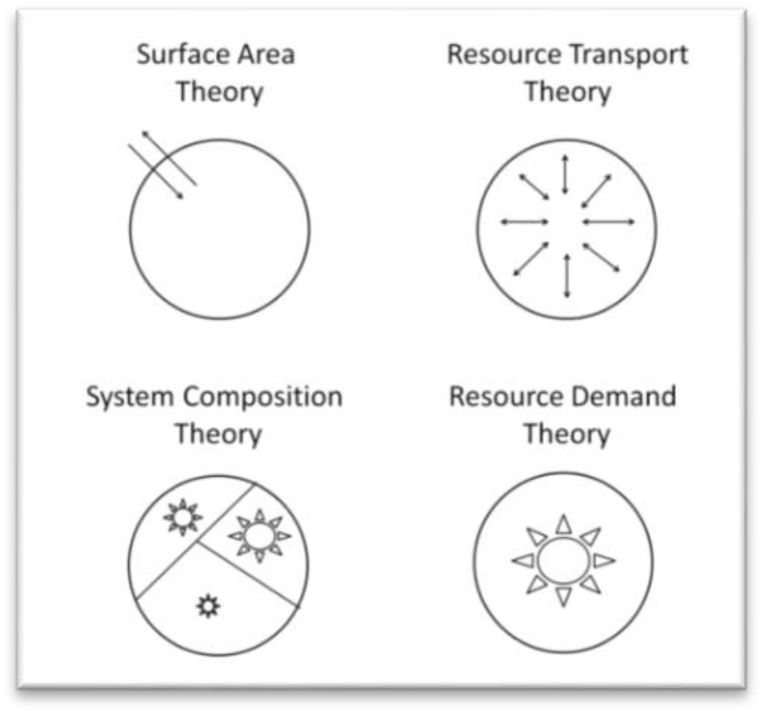
Schematic representations of the four major types of metabolic scaling theory: surface area (SA), resource transport (RT), system composition (SC) and resource demand (RD) theory. Cited from Glazier^26^.

As far as this article was concerned, the problem and principle summarized and proposed by Glazier^26^ were actually considered and followed in the dimension evolution analysis. For example, the evolutionary preference for relative surface area in prokaryotes and protist, the internalization of nutritional environment and therefore the dependence of internal resource transportation on the physical space dimension for animals, the coevolution of multiple metabolic links of fungi and the systematic composition of metabolic dimensions, and the constraints of the spatial nature of light (and water, nutrition) distribution on the evolutionary selection of metabolic dimension for plants, all of which were that life in evolution utilized the physical space dimensions and created biological dimensions by diversified approaches in metabolic scaling.

#### Unification of metabolic scaling theory in macroevolution

The major challenge of metabolic scaling research was the lack of a unified theoretical framework to summarize and sort out the current excessive diversity of theoretical assumptions and empirical research. As Bejan^47^ pointed out, the metabolic scaling law at the level of species classification just was the comprehensive result of multiple processes and causes, which was the inevitable problem to summarize the metabolic scaling law, and it was also the biggest logical misunderstanding to discuss the uniqueness of *b* value across species.

Glazier^17, 26^ had proposed that we can try to unify the theoretical research of metabolic proportion under the physical framework, and CMT framework undoubtedly pointed out a very correct direction. However, we still lacked a unified basis (that is, where does *b* come from?) and a unified benchmark (that is, where does *b* go?). Based on the analysis of the full text and the integration of relevant literature, it was believed that the golden section model of macroevolution provided such a unified theory framework, which defined the logic of dimension evolution of *b* as its source and the Fibonacci functional relationship of *b* as its distribution.

Similarly, metabolic homeostasis hypothesis^1^ and allometry metabolic scaling hypothesis^2^, can more play the role of metabolic scaling laws across species. The golden section law further became a rule to restrict the metabolic homeostasis law and allometry metabolic scaling law.

#### Another possible way based on biological time

Minimum Total Entropy theory^10^ believed that the value of *b* depends on the characteristic value of biological time (such as 1/4 of animals), but there was no theoretical support for the selection of biological time of different life forms in macroevolution.

In fact (or very likely), the dimensional evolution from one-dimensional growth to five-dimensional growth, had revealed a potential evolutionary logic of biological time in macroevolution: biological times might just be the reciprocal of dimension application of different life forms. However, the most important question was, what was the exact connotation of biological time in macroevolution? A rhythm related to metabolic macroevolution under the constraint of nonequilibrium thermodynamic system? This was an interesting question.

### A1.4 Macroevolutionary geometry and dimensional evolution

So far, the geometric operations in dimensional evolution has not really involved such methods as fractal geometry. It was argued that in macroevolution, the emergence of fractal growth^8^ might only be a skill of spatial utilization of biology, rather than the geometric basis that determines the logic of dimensional evolution. On the contrary, the least carrier network theory^30^, which did not depend on fractal networks, showed the applicability of a more general dimensional evolution model for animals.

In terms of macroevolution, the geometric analysis method of biological dimension should be different from the current overly refined and microscopic analysis paradigm in metabolic scaling. Of course, the combination of spatial geometric analysis and physical-chemical mechanism models in metabolic processes was still the magic weapon to reveal the diversity of metabolic scaling law in different levels of evolution.

Based on the experience of dimension analysis in macroevolution in this paper, it was believed that a new category should be considered: macroevolutionary geometry. Different from metabolic scaling studies and paleontology, its basic feature in geometric analysis was "*thick lines, large outlines*", so as to better grasp the macroevolutionary trend of biological morphological structure, reveal the dependent evolution trajectory of its metabolic function, and find the macro logic of cross-boundary evolution. At the same time, for the dimensional analysis in the secondary evolution, priority should be given to the highest level classification system, to extract the main thread of dimensional evolution from the complex natural selection.

The spatial dimension analysis of biology does not need to accurately measure its length, area and volume, but focuses on establishing the dimension size and geometric operation relationship of length, area and volume within the scope of biological activities^5^.

#### Technical principle_Growth principle

It has always been emphasized "spatial growth mode", rather than "spatial mode". If it was only statically applied the geometric operation logic of physical space in evolution, it will bring many problems. For example, in the two-dimensional growth of protist, although the surface area is still an important metabolic limiting link, this evolution trend will lead to the synergy of multiple metabolic sites within cells rising to the primary metabolic limiting factor following the growth of individuals. If the general relation between area and metabolism in physical space still is mechanically applied, obviously, the metabolic dimension generation mode of protist in evolution will be omitted.

*B-M* is not a static relationship, but a dynamic one, emphasizing the distribution trend of metabolic and non-metabolic parts in growth and evolution. In practical application, the geometric relationship of biological volume, area and length will deviate from Euclidean geometric logic due to the change of growth mode, which is a basic principle of geometric operation in evolution.

This principle also could be applied in other levels of species classification, even including the comparison of different stages of individual life history. For example, the spatial growth patterns of young trees and big trees are completely different, so it also applies to different geometry rules of dimensional generation. This principle is called the growth principle. In practical analysis, the growth principle of and the relative principle (in following) can be cross applied.

#### Technical principle_Extreme principle

The strict adherence of macroevolution to golden section often leads to the "mechanized" borrowing of some basic physical laws by biological individuals. For example, the morphological evolution of tall trees borrows from the geometric laws of tree mechanics, and the internal circulation system evolution of mammals borrows from fractal geometry. As some skills, we can use some physical mechanics and solid geometry calculation formulas to assist in the construction of macroevolutionary geometry model. For example, hexahedral bacteriophages have the inevitability and rationality of their mechanical individual structure evolution under the macroevolutionary logic (appendix one-A1.5). For this life form, the application of some pure physical routes in dimensional analysis is inseparable. It is called the extreme principle.

#### Technical principle_Trending principle

The primary purpose of dimensional analysis in macroevolution is to find the trend and direction of evolution. We may not find strictly corresponding species records in fossil evidence during the construction of geometric models, but as long as this trend of dimensional evolution has great evolutionary significance and indication, we can accept the results from geometric model in principle. For example, we speculate that there should be a two-dimensional early prokaryote (or archaea) group in early evolution stage, but we can’t find this clue in fossil evidence and prokaryote remains, but the two-dimensional dimensional evolutionary logic and its great macroevolutionary significance are beyond doubt. It is called the trending principle.

#### Technical principle_Relative principle

For example, the relative two-dimensional concept of resource distribution mentioned in the macroevolution of plant absorption mode for light, nutrients and water. However, this relativity should be not suitable in secondary evolution or microevolution. To some extent, species differentiation is the niche segmentation of subtle differences in the spatial distribution of resources, such as shallow root plants and deep root ones. However, in macroevolution, we do not need to consider the specific temporal and spatial changes of resources too much, but to start with the essence of the spatial supply mode of resources and find the key factors in macroevolution. Similarly, this principle is also applicable to the relative filling process of fungi to three-dimensional nutrients, as well as the "spatial internalization" mode of animals. Because strictly speaking, the filling of three-dimensional space is relative whether for early animals or fungi, compared with the standard of fractal network in higher animals. However, this does not change the geometric essence of their dimension application in macroevolution. It is called the relative principle.

### A1.5 Fibonacci code box in macroevolution

#### Position of hexahedral phage in macroevolution (n=6)

The basic structure of virus was a layer of protein shell wrapped in a mass of nucleic acid material, and its basic metabolism (activity) was to invade the interior of the host cell to complete replication and proliferation. The primitive virus in general had a coat shell, which was of shape as an irregular cake. From the perspective of evolution, it was not conducive to entry or release. Therefore, the morphology gradually evolved into the form of hexahedron, which had the advantages of stable structure (to protect nucleic acid substances). At the same time, special adsorption (six legs of bacteriophages) and release structures also evolved (the bulge at the top of the prism should be related to release function to breaking wall of host cell). Therefore, the six growth dimensions of hexahedron bacteriophage included 3 of hexahedron (three dimensional physical structure), 1 of release structure (one dimensional physical structure) and 2 of adsorption structure (related with the host cell surface area).

The only purpose of viral metabolism (activity) just was to replicate by borrowing bodies. The complete links of virus replication (metabolism) include adsorption (determined by cell surface area, 2-dimensional), entry (related to distance, 1-dimensional), replication (related to distance, 1-dimensional), assembly (related to distance, 1-dimensional), release (using the cell metabolic system to complete proliferation and growth before release, related to cell volume, 3-dimensional), from which eight metabolic dimensions therefore can be formed. Five dimension were related to the host cell structure, the other three were the special needs of viruses. Thus, for hexahedral bacteriophage *D_M_*=6, *D_B_*=8, *b*=8/6=4/3.

#### Position of nervous system in macroevolution (n=7)

In 2017, Reimann et al.^48^ found that our brain was full of multidimensional geometric structures and even operated in 11 dimensions. For the position of *n*=7, nervous system was worth looking forward to. We speculated that the evolution of the nervous system was also difficult to escape the constraints of the golden section law, and its metabolic dimension (i.e. working dimension) selected in macroevolution should be 13, not just 11.

First of all, different from all the previous life forms, the nervous system was a non-independent subsystem of life. The evolutionary significance of its existence was information processing (that was, the metabolic activities of the nervous system) to coordinate the overall activities of individual life. Therefore, first of all, there were five growth dimensions of nervous system directly come from individual life: nervous system should be distributed throughout the body, including three dimensions of the body and two ones from the body surface. Different from mycelium, the extra two come from the structure of dendrites. The remaining two dimensions should come from the cerebral cortex, which was a two-dimensional structure, so the growth dimension number of nervous system was 7.

The complete working mode of the nervous system required different types of neurons (including afferent ones∝*L*^3+2^, efferent∝*L*^3+2^, processing∝*L*^2+1^, *L* biological length) to coordinate and control different activities at the same time. Therefore, the generation of metabolic dimension was actually the multiplication of three types of neuronal activities (*L*^5^ * *L*^5^ * *L*^3^). The dimensions of afferent and efferent actions corresponded to five dimensions of growth respectively (three dimensions came from the radial distribution of axons in the body, two dimensions came from the distribution of dendrites at the end of body), and three dimensions of processing action was the three-dimensional packing in the folded cortex of the dendritic interwoven network). Therefore, the metabolic dimension number of nervous system was 13. Thus, for nervous system in macroevolution, *D_M_*=7, *D_B_*=13, *b*=13/7.

#### Fibonacci code box of life evolution

Thus, the first seven of the Fibonacci series (excluding *f* (0)) was likely to be such an order from prokaryotes, protist, plants, animals, fungi, viruses, to nervous system (Table 5 and Fig.7). The last one was left to the nervous system, should be very interesting. The emergence of wisdom was a functional by-product of life evolution, as a management organization with the evolution from simple to complex.

**Table 5.**
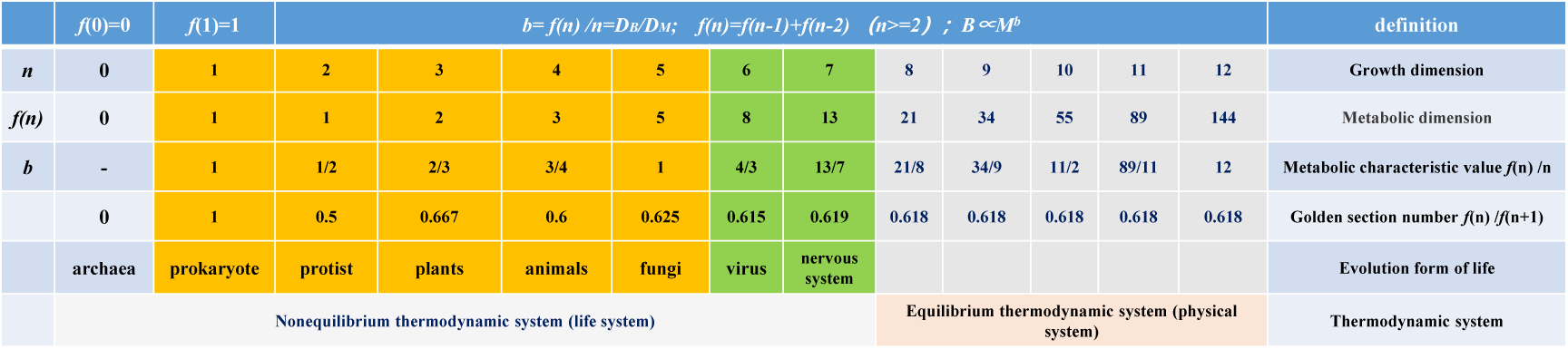
Fibonacci code box of life evolution.

**Fig. 7.**
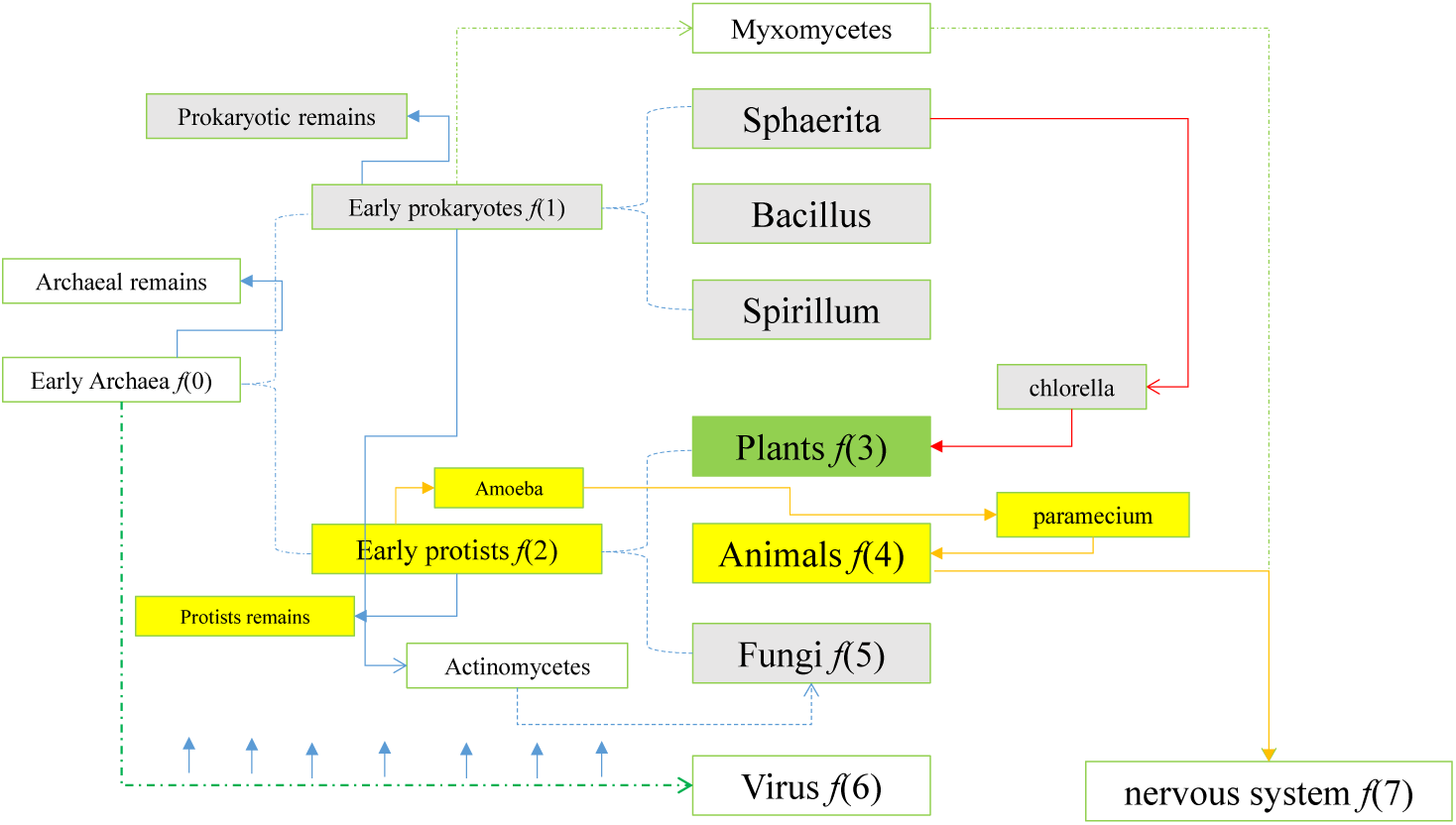
Another tree of life evolution.

In the existing fossil records, only archaea had life forms with regular geometric. From the logic of morphological evolution and dimensional evolution, it was easy to associate the origin of viruses. Myxobacteria was the only organism with potential memory ability found in lower life forms, which was also the space we can imagine.

## Appendix two. Geometric derivation of dimension application and metabolic realization in animals

The least carrier theory^30^ explained the general logic and basic mode of animal dimension evolution in macroevolution, compared with WBE model^8^. Based on this model, this paper attempted to build a unified geometric model of dimension application and metabolism realization in animals.

### Geometric prototype of metabolic scaling in the least carrier network theory^30^ (compiled with reference to the summary of Han and Fang^6^)

According to the least carrier network theory^30^, the raw materials required for animal metabolism were transported from a supply center to various metabolic sites through the network. In order to receive nutrition everywhere, this site must be filled with the whole body. The metabolic rate in each site was not related with body size. Therefore, the total number of metabolic sites was *L^D^*, where the length variable *L* was the unit of the average distance between sites, and *D* was the Euclidean space dimension filled by the network system. Since every place needs to receive nutrition, the metabolic rate is

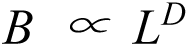

Substances needed to be transmitted to each site by carriers (such as blood), so the network needed to be filled with a certain volume of carriers. The volume of carriers *C* depended on the network structure. Banavar et al.^30^ believed that the higher the transmission efficiency, the less the capacity of the carrier required by the network.

Therefore, according to the law of mass conservation, the allometric relationship of *C* can be derived,

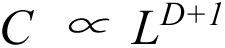

There was,

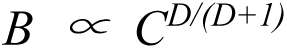

And because *C* was isometric to the size of the organism *M*, that was, *C ∝ M.* Therefore, the scaling relationship between metabolic rate *B* and individual size *M* was,

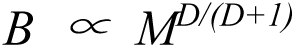

### "spatial internalization" and animal basic metabolic dimension D_BV_

According to the geometric prototype of the least carrier network theory^30^, whether for lower animals or higher animals, the spatial growth parameters that limit metabolic efficiency, including the average distance *L* between metabolic sites scattered throughout the body, and the Euclidean space dimension *D* filled by the network system, were limited by this "center-diffusion" mode of metabolic dimension generation, and therefore the geometric operation relationship would not change.

This was the geometric essence of the metabolic evolution model of "spatial internalization" for animals, This metabolic dimension, we called it as the basic metabolic dimension of animals *D_BV_*=*D*=3.

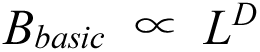

### Dimensional application and metabolic realization of mammals

The essence of metabolic evolution just was to "obtain the maximum metabolic benefit at the minimum cost of growth". Banavar et al.^30^ believed that the higher the transmission efficiency, the less the capacity of the carrier (*C*) required by the network, which was fully consistent with the essence of metabolic evolution. From *C∝M∝ L^D+^*^1^, the growth dimension *D_M_*=*D*+1=4. Therefore, for mammals that have evolved a perfect internal circulation system, there were,

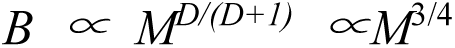

### Dimension application and metabolic realization of protozoon and ectotherm

For protozoon and ectotherm that still relied on the body surface for respiration and temperature regulation, the limitation of body surface on metabolism could not be ignored. Therefore, a metabolic dimension determined by body surface area was added. Similarly, according to the basic principles of metabolic evolution and mass conservation, this increased body-surface metabolic dimension *D_BS_* was equal to the growth dimension *D_M_* minus the basic metabolic dimension *D_BV_*, that was,

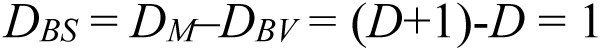

Therefore the total metabolic dimension *D_B_* = *D_BS_* + *D_BV_ =D*+1= *D_M_*. For protozoon and ectotherm there was,

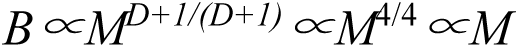

First, there were two contradictory demands for body surface temperature control of variable temperature animals. One was to receive enough sunlight radiation (the larger the area, the better), and the other was to control the rapid loss of body temperature (the smaller the area, the better). Therefore, in evolution, animals generally chose elongated body structures, one was to increase the relative surface area (improve the relative diffusion efficiency) and the other was to reduce the absolute surface area (reduce the absolute diffusion rate). At the same time, affected by the solar radiation mode, only one side of the animal can receive radiation at the same time. Therefore, according to the general principle of macroevolutionary geometry analysis, it can be deduced that the key geometric parameter limiting the metabolic efficiency of the body surface was the body length L, rather than the absolute surface area. Therefore, for variable temperature animals, there was *A_BS_*∝*L*, *D_BS_*=1. In fact, with the evolution of animals, various auxiliary measures have evolved for the control of body surface temperature, such as body hair, scales, fat layer, etc., which further improved the effect of animals on the control of body surface temperature, gradually got rid of absolute dependence on the geometric parameters of body surface area.

Secondly, for unicellular protozoon such as Paramecium, body surface metabolism (respiration) was mainly subordinate to the metabolic spatial growth mode of "spatial internalization". The coelenteratic system of Paramecium was relatively developed (from mouth to abdomen, throughout the body), due to the lack of a complete internal circulation system, to improve the internal nutrition supply efficiency of "center-radiation" (shorten the internal transmission distance and coordinate with the body surface metabolism). Therefore, the key geometric parameter limiting body surface metabolism, body surface area, also depended on the spatial growth mode of the coelenteratic system, which was obviously a one-dimensional growth mode.

Therefore, Paramecium still exists *A_BS_*∝*L*, *D_BS_*=1. The mode could also be considered as a transformation of geometric relationship caused by the change of spatial growth mode for the dimensional relationship between mitochondrial distribution and cell volume^21, 22^.

### Unified geometric model of dimensional evolution and metabolic realization in animal kingdom

However, for mammals, the limiting effect of body surface on metabolism had been lost, therefore *D_BS_*=0. But as the growth cost of metabolic evolution, this dimension still existed in *D_M_*.

Therefore, a unified metabolic scaling relationship in animals could be described as,

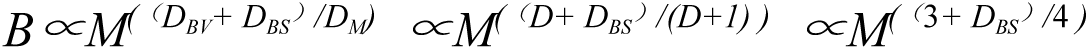

Among them, *D_BV_* is the basic metabolic dimension, *D_BS_* is the surface metabolic dimension, *D* is the physical spatial dimension occupied by individuals, which is equal to 3, and *D_BV_=D*.*D_BS_*=1 for low and variable temperature animals, and *D_BS_*=0 for constant temperature animals with fully developed internal circulation.

## Appendix three. Geometric derivation of dimension application and metabolic realization in plants

Unlike animals, there was no unified geometric model for plants just for their high morphological plasticity. For higher plants, the premise of geometric derivation was that there was strictly light restriction on metabolism, with complete differentiation of leaves and vascular organs.

### General Euclidean geometric operation rules of higher plants

For higher plants strictly following the three-dimensional growth based on Euclidean geometry, there was

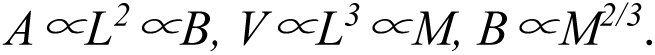

Plants were typical representatives of three-dimensional growth in evolution. From sphaerita to chlorella, mosses, small grass and high trees, they might all follow the three-dimensional conservative evolutionary route. However, for early aquatic photosynthetic bacteria and unicellular photosynthetic algae, their photosynthetic metabolic organs were mainly chlorophyll macromolecules filled in the globular cells, rather than in the cell surface, to get higher metabolic rates, therefore *b*=*D_B_/D_M_*=3/3=1 can be deduced when metabolic rates were strictly limited by cell volume^13^. Only after higher terrestrial plants evolved leaves and vascular bundles, the geometric spatial behavior of growth and metabolism separated, and finally evolved to the 2/3 rule.

### Tall and mature trees

By another way, there is *V*∝*D*^2^*H*∝*M* for trees in general, *V*, tree volume, *H*, tree height, *D*, trunk diameter. It could be easily deduced *V*∝*H^4^*∝*M* when^28, 29^ *H*∝*D*^2/3^, just as the 3/4 rule from WBE model^40^. However, the growth state of tall and mature trees must be considered, which will lead to the change of the geometric relationship between *H* and *D*. The height growth of tall and mature tree is very slow in general^49, 50^. Therefore, relative to *D*, *H* is approximately constant in the growth. So there is approximately *V*∝*D*^2^ for the tall and mature trees, rather than *V*∝*D*^2^*H*. The result is as same as that in the method. For *H*∝*D*^2/3^(To a great extent, the relation only is true in such a critical state), *V*∝*H*^3^∝*L*^3^ can be deduced in some critical state. Therefore, the applying of *H*∝*D*^2/3^ must be limited, otherwise the same questions (or debates) would happen repeatedly^9, 51^. Just from the comparing between adult trees and young ones, the problem could be seen. The difference of dimension analysis in metabolic scaling between in evolution and in biology should be distinguished clearly, which is very important.

In addition, there is total leaf mass *R*∝canopy volume *V_leaf_* ∝*r*^2^*h*, *h* crown thickness (nearly a constant in some critical state), *r,* crown radius. It can be deduced that *R*∝*r^2^,* for *R*∝*B,* so *B*∝*r^2^*. According to empirical research results^52^, the Hurst exponent^53^ characterizing the scaling of the crown radius (*r*) with height (*H*), there was *r*∝*H* in in tropical forests, which could be consider as the represent of tall and mature trees. So there is *R*∝*B*∝*r^2^*∝*H*^2^∝*L*^2^, *D_B_*=2, *b*= *D_B_* /*D_M_* =2/3.

### Young tree individuals living in initial dense group

For young tree individuals living in initial dense groups, the mechanical limit disappeared, that was, *H*∝*D*^2/3^ was no longer established. From *V*∝*D*^2^*H*, because the tree body was completely dominated by high growth, *D* of young trees was approximately constant in the growth compared with *H*, so there was *V*∝*H*∝*L*∝*M*, *D_M_*=1 can be deduced.

For the *D_B_*, it could be deduced that the projected area of each young tree individual was unchanged in an initial dense group (radius of canopy *r* was a constant), there was total leaf mass *R*∝ canopy volume *V_leaf_* ∝*r*^2^*h*, *H*∝*h, h*, the length of canopy. For *r* was a constant, it could be deduced that *R*∝*H,* for *R*∝*B,* so *B* ∝*H*∝*L*∝*M*, *D_B_*=1, *b*= *D_B_* /*D_M_* =1. The result was completely consistent with Reich et al^51^.

### Self-thinning of plant population

Among higher life forms, group living was the most common form for plants. This form (selection) in evolution actually existed from the beginning of photosynthetic bacteria. Group living was a sufficient condition for some higher plants to quickly achieve the 2/3 rule, which was also an important condition for higher plants to achieve energy equivalence after the evolution of complete leaf crowns and vascular bundles. The direct result of energy equivalence was *N*B* ∝ *M*^0^ (population productivity was constant), and there was *N*∝*M*^-3/2^ along self-thinning trajectory^15^, *B* ∝*M*^2/3^ therefore could be derived. When *B*∝*M*, as long as the energy equivalence was met, *N*M*∝*M*^0^ (population biomass was constant) could also be derived. These were two manifestations of constant output law. The latter could be found in asexual reed population and annual grass population.

### Is plant suitable for 3/4 in macroevolution? Counter evidence of one typical research case

Enquist et al.^9^ and West et al.^40^ constructed 3/4 metabolic model of plants. According to the model, there were *Q*∝*D*^2^, *D*∝*M*^3/8^, so *Q*∝*M*^3/4^∝*B*. And it was very close to the empirical value *D*∝*M*^0.412^. *Q*, xylem sap flow rate (*v/d*), *D*, trunk diameter. Similarly, we can also draw the following conclusions based on some of their derivation.

First of all, *Q* could be easily converted equivalently to the total xylem sap *Q_M_* in a day, and the scaling relationship between *Q*_M_ and *D* would not be changed (*Q*_M_∝*D*^2^). According to the thought of least carrier theory^30^, *Q*_M_∝*M*∝*D*^2^∝*Q* can be deduced just as that in animals, instead of *Q*∝*D*^2^∝*B*. At least in terms of evolution, we believed that the least carrier theory revealed the basic principles of metabolic evolution, “*obtain the maximum metabolic benefit at the minimum cost of growth*”.

Secondly, *D*∝*M*^1/2^ was also not far from *D*∝*M*^0.412^. And logically, the theoretical value should be higher than 0.412 (the empirical value *D*∝*M*^0.412^). In terms of the metabolism scaling law for tree in evolution, no matter 3/4 or 2/3 rule, the individual object would not be a young tree in complete height growing (for it *D* ∝ *M*^0^), but an adult tree following basic mechanical constraints (namely *H*∝*D*^2/3^). However, in practices it generally was difficult to reach the theoretical growth stage for the invested trees. Let’s make a simple sort of 0, 3/8, 0.412, 1/2. Obviously, 1/2 was closer to the theoretical value.

As before, *B*∝*M*^2/3^ can be deduced from *H*∝*D*^2/3^. Therefore, the metabolic scaling of adult and tall trees in macroevolution, strictly followed the geometric operation rules of Euclidean space.

